# Machine learning reveals the control mechanics of an insect wing hinge

**DOI:** 10.1101/2023.06.29.547116

**Authors:** Johan M. Melis, Igor Siwanowicz, Michael H. Dickinson

## Abstract

Insects constitute the most species-rich radiation of metazoa, a success due to the evolution of active flight. Unlike pterosaurs, birds, and bats, the wings of insects did not evolve from legs^1^, but are novel structures attached to the body via a biomechanically complex hinge that transforms tiny, high-frequency oscillations of specialized power muscles into the sweeping back-and-forth motion of the wings^2^. The hinge consists of a system of tiny, hardened structures called sclerites that are interconnected to one another via flexible joints and regulated by the activity of specialized control muscles. Here, we imaged the activity of these muscles in a fly using a genetically encoded calcium indicator, while simultaneously tracking the 3D motion of the wings with high-speed cameras. Using machine learning approaches, we created a convolutional neural network^3^ that accurately predicts wing motion from the activity of the steering muscles, and an encoder-decoder^4^ that predicts the role of the individual sclerites on wing motion. By replaying patterns of wing motion on a dynamically scaled robotic fly, we quantified the effects of steering muscle activity on aerodynamic forces. A physics-based simulation that incorporates our model of the hinge generates flight maneuvers that are remarkably similar to those of free flying flies. This integrative, multi-disciplinary approach reveals the mechanical control logic of the insect wing hinge, arguably among the most sophisticated and evolutionarily important skeletal structures in the natural world.

---

Whether to forage, migrate, reproduce, or avoid predators, aerial maneuverability is essential for flying insects. Most insects actuate their wings using two morphologically and functionally distinct sets of muscles. Contractions of the large, indirect flight muscles (IFMs) are activated mechanically by stretch rather than by individual action potentials in their motor neurons—a specialization that permits the production of elevated power at high wingbeat frequency^5–7^. There are two sets of IFMs, dorso-ventral muscles (DVMs) and the dorso-longitudinal muscles (DLMs), arranged orthogonally to maintain a self-sustaining oscillation (Fig. 1a-c). The tiny deformations in the exoskeleton generated by the IFMs are mechanically amplified by the wing hinge^8–12^, an intricate structure consisting of an interconnected set of small sclerites embedded within more flexible exoskeleton and regulated by a set of small control muscles^13^ (Fig. 1a,b). In this paper, we use machine learning approaches and physics-based simulations to gain insight into the underlying mechanics of the wing hinge of flies and its active regulation during flight.

**Figure 1.**
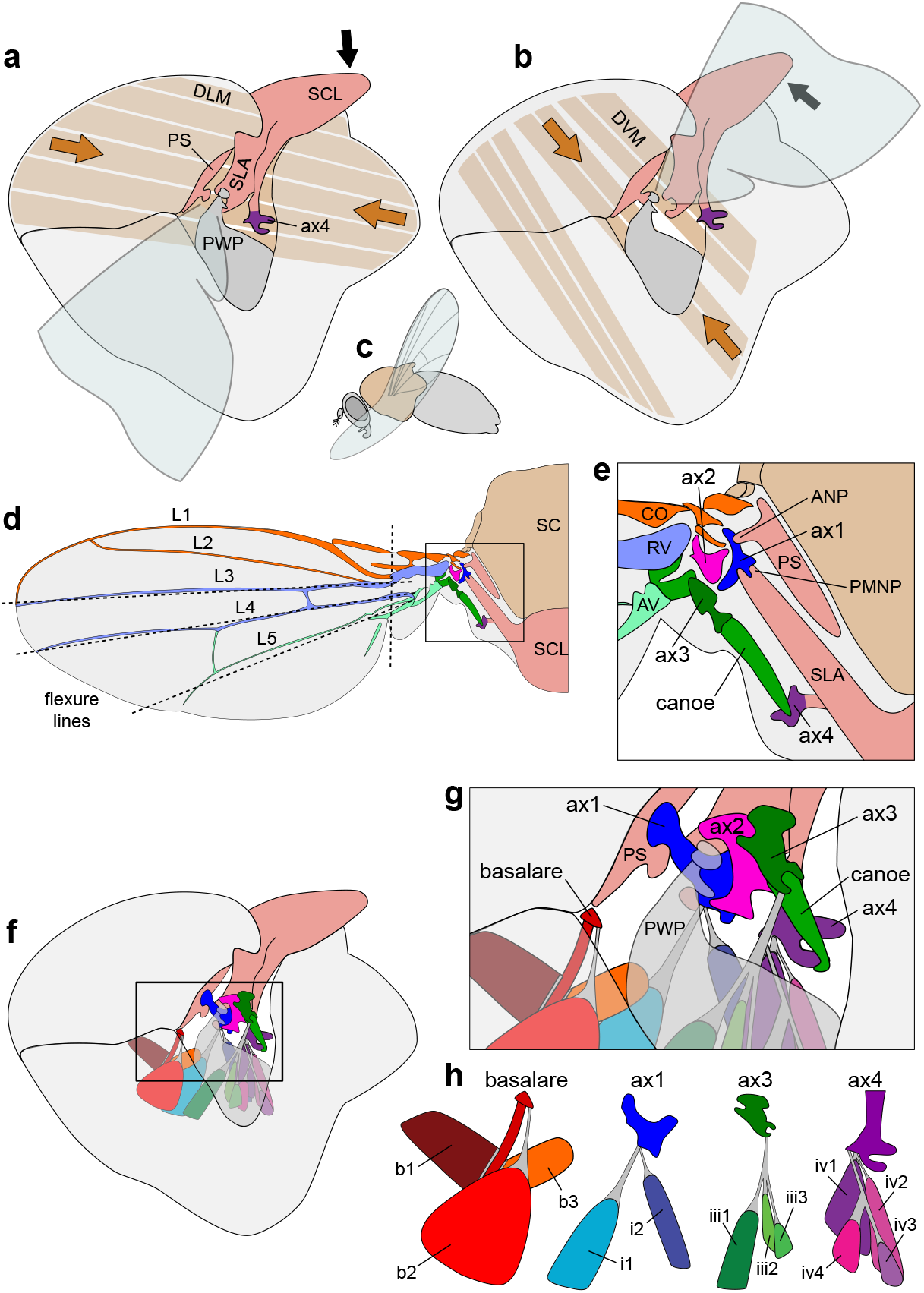
The wing hinge of a fly is actuated by large power muscles and controlled by small steering muscles. The schematic drawings are based on a synthesis of prior work^2,7-18^ and our own new morphological analysis (Supplementary Video 1). **a**, Contraction of the dorso-longitudinal muscles (DLMs) causes the scutellum (SCL) to move downward. **b**, Contraction of the dorso-ventral muscles (DVMs) cause the SCL to move upward. These motions are transferred to the wing hinge via the PS and SLA, with the wing’s fulcrum located at the pleural wing process (PWP). **c**, Side view of a flying fly indicating orientation in panels (a) and (b). **d**, Schematic top view of the thorax, hinge, and wing, illustrating wing veins (L1-5), scutum (SC), and scutellum (SCL). Dotted lines represent hypothesized flexure lines on the wing. **e**, Close-up of the wing hinge highlighting the parascutal shelf (PS), anterior notal process (ANP), scutellar lever arm (SLA), posterior-medial notal process (PMNP), first, second, third, and fourth axillary sclerites (ax1, ax2, ax3, ax4), costal vein (CO), radial vein (RV), anal vein (AV), and a previously unnamed we term the canoe. **f**, Layout of the steering muscles and sclerites in the thorax. **g**, Close-up view of the basalare, axillary sclerites, and steering muscles. Tendons are marked in gray. **h**, Steering muscles grouped according to the sclerite to which they attached: basalare (b1, b2, b3), first axillary (i1, i2), third axillary (iii1, iii2, ii3), and fourth axillary (iv1, iv2, iv3, iv4) muscles.

## Morphology of the dipteran wing hinge

The 3-dimensional structure of the wing hinge of flies is so complex that it is difficult to depict in 2-dimentional drawings. As an introduction to this intricate structure, we provide an animation in Supplementary Movie 1 that illustrates the arrangement of wing sclerites within the hinge and the control muscles that regulate their function. These data are based on our ongoing effort to accurately reconstruct the detailed morphology of the wing hinge of fruit flies (*Drosophila melanogaster*), using a specialized application of confocal microscopy. While this efforts provides some clarity on the morphology of the hinge in a static configuration—with the wings folded snuggly against the body—the mechanical operation of the hinge during flight remains enigmatic, because the wing sclerites are difficult to visualize externally and move so rapidly that their changing configuration has not been accurately captured by either stroboscopic photography^14^, high speed videography^15^, or X-ray tomography^16^. The consensus of all prior morphological analyses in various dipteran species^2,9,10,12,16^ suggests that the tiny deformations of the thorax generated by contractions of the IFMs are transformed into wing motion via the mechanical actions of the anterior and posterior-medial notal wing processes (ANP and PMNP) on the ‘x’-shaped 1^st^ axillary sclerite (ax1) at the base of the wing (Fig. 1d-h). The ANP projects from the anterior end of the parascutal shelf and fits in the groove between the two anterior lobes of ax1. The PMNP sits at the anterior end of the large scutellar lever arm (SLA) in a notch between the two ventral lobes of ax1. Contractions of the DLMs during the downstroke are thought to be transmitted to the wing hinge via two main mechanisms: (1) a rotation of the scutellum along its junction with the notum that moves the scutellar lever arm and PMNP upward, and (2) an elevation of the entire notum that raises the ANP at the lateral edge of the parascutal shelf (Fig. 1a). Contractions of the antagonistic DVMs during the upstroke cause the opposite effects (Fig. 1b). The more distal 2^nd^ axillary sclerite (ax2) possesses a dorsal flange that fits between the two dorsal protuberances of ax1, coupling the two sclerites during flight (Fig. 1g, Supplementary Movie 1). Ax2 is a critical component of the hinge system because it is rigidly attached to the radial vein—the main structural spar of the wing, and it has an anteriorly projecting ventral extension that fits into a notch in the plural wing process (PWP), thus forming the main fulcrum for wing motion. Because this fulcrum point is distal and anterior to ax1, upward motion of the SLA causes downward motion and pronation of the wing, whereas downward motion of the SLA causes upward motion and supination of the wing^8^.

In addition to the SLA, the anterior end of the scutellum is connected to another anteriorly projecting structure, the 4^th^ axillary sclerite (ax4), providing a second mechanical pathway linking IFM strains to the wing base (Fig. 1a-e). Ax4 lies ventrolateral to the SLA and is attached by a long, previously unnamed sclerite (which we term the ‘canoe’ based on its peculiar shape) to the 3^rd^ axillary sclerite (ax3). Ax3 contacts ax1, ax2, and the base of the anal vein, which serves as the main structural spar for the posterior half of the wing (Fig. 1d,e). Due to its connection with the anal vein, the linkage system consisting of the scutellum, ax4, canoe, and ax3 could influence wing camber and angle of attack during flight^10,17^, but this functional role remains speculative. We note that some prior authors have drawn the canoe as an extension of ax3^17,18^, but the two structures are clearly distinct in our confocal reconstructions (Supplementary Movie 1) and more recent μ-CT data^19^.

In addition to the four axillary sclerites that are directly incorporated into the hinge, another important sclerite, the basalare, sits in the membranous episternal cleft just anterior to the wing (Fig. 1g,h)^14,16,20^. From the surface, the basalare is a triangular-shaped structure, but it has a long internal invagination (apophysis) that provides an attachment site for two of its three control muscles (Fig. 1h). During flight, the basalare is thought to oscillate passively due to the opening and closing of the episternal cleft and via its ligamentous connection to the base the radial vein. However, the magnitude of these oscillations and the mean position of the basalare is actively regulated by its three control muscles^16,21^, thereby adjusting the tension of the ligamentous connection to the radial vein.

Although the IFMs provide the power to oscillate the wings, they cannot regulate the rapid changes in kinematics required for flight maneuvers or the sustained asymmetrical patterns of motion necessary to remain aloft with damaged wings^22^. In flies, that control is mediated by a set of 12 steering muscles that insert on three of the four sclerites (ax1, ax3, ax4) and the basalare (Fig. 1f-h, Supplementary Movie 1). The steering muscles are named after the sclerite on which they act (ax1: i1 and i2; ax3: iii1, iii2, iii3; ax4: iv1, iv2, iv3, and iv4; basalare: b1, b2, and b3). Ax4 and its muscles are sometimes described using an alternative nomenclature, hg, after its German name ‘hinter-Gelenkforsatz’^18^. Each steering muscle is innervated by only a single excitatory motor neuron^23,24^, thus the entire flight motor system is remarkably sparse relative to comparably maneuverable vertebrates such as hummingbirds^25^.

## Neural network model wing hinge

Machine learning is proving increasing useful at gaining insight into many phenomena in biology, for example, the processing of visual information by mammalian cortex^26^. While such approaches are powerful in their ability to accurately predict an output (e.g. response of a neuron) from a set of inputs (e.g. complex sensory stimuli), it remains a challenge to gain mechanistic insight from the hidden features of a neural network after its successful training. Our aim is to use machine learning to probe the musculoskeletal transformations that underlie the mechanical function of the hinge, using this activity of control muscles as an input, and the detailed 3-D motion of the wings as an output. We constructed an experimental apparatus that allowed us to capture the wing motion of tethered flies at 15,000 frames per second with three high-speed cameras, while simultaneously measuring steering muscle activity using the Ca^2+^ indicator, GCaMP7f^27^ (Fig. 2, Extended Data Fig. 1). To capture data that represented a wide range of muscle activity, we collected a large set of sequences, each triggered whenever the activity of one of the steering muscles (chosen *a priori* at the start of each trial) exceeded a user-defined threshold. To encourage a wide variety of flight behaviors, we also displayed open- and closed-loop visual stimuli on a cylindrical array of LEDs^28^ surrounding the fly. In total, we recorded a total of 485 flight sequences from 82 flies. After excluding a subset of wingbeats from sequences when the fly either stopped flying or flew at an abnormally low wingbeat frequency, we obtained a final dataset of 72,219 wingbeats.

**Figure 2.**
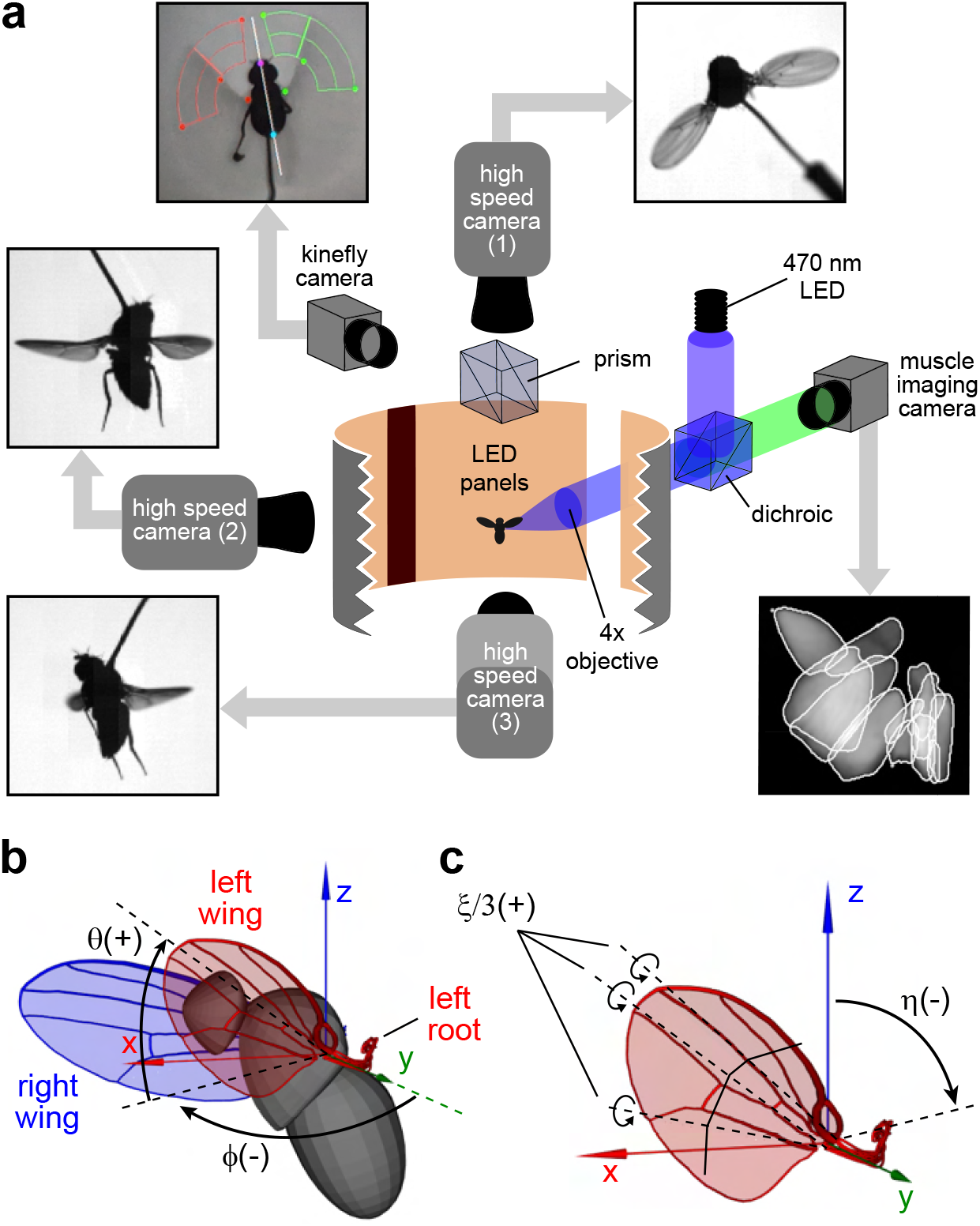
Simultaneous imaging of muscle activity and wing motion. **a**, Three high-speed cameras with IR-backlighting capture a tethered fly from three orthogonal angles. An epifluorescence microscope projects blue light onto the left side of the fly’s thorax for imaging muscle activity using GCaMP7f. Visual stimuli are provided by an array of orange LED panels surrounding the fly. A prism splits the top view between a high-speed camera and a kinefly camera (see online methods) that tracked the stroke amplitude of the two wings for use in epochs of closed-loop control of the visual display. **b**, Wing pose is determined from the high-speed camera footage using custom machine vision software. The wing state is subsequently described by four angles relative to the Strokeplane Reference Frame (SRF). The SRF is established by performing principal component analysis (PCA) on the motion of the left and right wing roots, with the x-y plane representing the mean strokeplane. Two angles, *ϕ* (stroke angle) and θ (deviation angle), describe the left wing’s orientation in the SRF (the +/-sign indicates the direction of rotation). **c**, The wing pitch angle, η, indicates the orientation of the leading edge relative to the z-axis of the SRF. The deformation angle, ξ, represents the uniform rotation along three hinge lines that roughly align with the L3, L4, and L5 veins.

To reconstruct the kinematics of the wings from the high-speed data, we developed an automated tracking algorithm that used a trained convolutional neural network (CNN)^29^ and particle swarm optimization^30^ (Extended Data Fig. 2a), a work flow that is described in detail in Supplementary Information. We specified wing pose relative to a fixed reference frame that approximated the mean stroke plane using three Tait-Bryan angles: stroke position (*ϕ*), deviation (*θ*), and wing pitch (*η*) (Fig. 2b). We also defined a fourth angle (*ξ*) that quantified the chord-wise deformation of the wing (i.e. camber) from the leading to the trailing edge (Fig. 2c). We parsed the wing kinematic traces into separate wingbeats, beginning and ending at dorsal stroke reversal. To reduce the dimensionality of the data, we fitted Legendre polynomials to all four kinematic angles, creating a vector of 80 coefficients (*ϕ* = 16, *θ* = 20, *η* = 24, *ξ* = 20) that accurately captures the motion of one wing during each stroke.

We determined the Ca^2+^ signals originating from each of the 12 steering muscles using a real-time unmixing model as previously described^27^. The muscle activity traces were normalized over each trial duration, such that 0 and 1 corresponded to -2 and +2 standard deviations, respectively (Fig. 3a). Due to its binding kinetics^31^, GCaMP7f reports a low-pass filtered version of sarcoplasmic Ca^2+^ levels, which in addition to attenuating high frequencies, introduces a time delay between changes in ion concentration and fluorescence signals. Because of these effects, the muscle signals relevant for the kinematics of any individual stroke are spread out over several stroke cycles, and the GCaMP7f data recorded after a particular wingbeat are better representative of the Ca^2+^ levels in the muscles at the time the kinematic measurements were made. Thus, when training and evaluating our CNN, we associated the kinematics of each individual wingbeat with the Ca^2+^ signals recorded in the muscles the during the subsequent 9 wingbeats.

**Figure 3.**
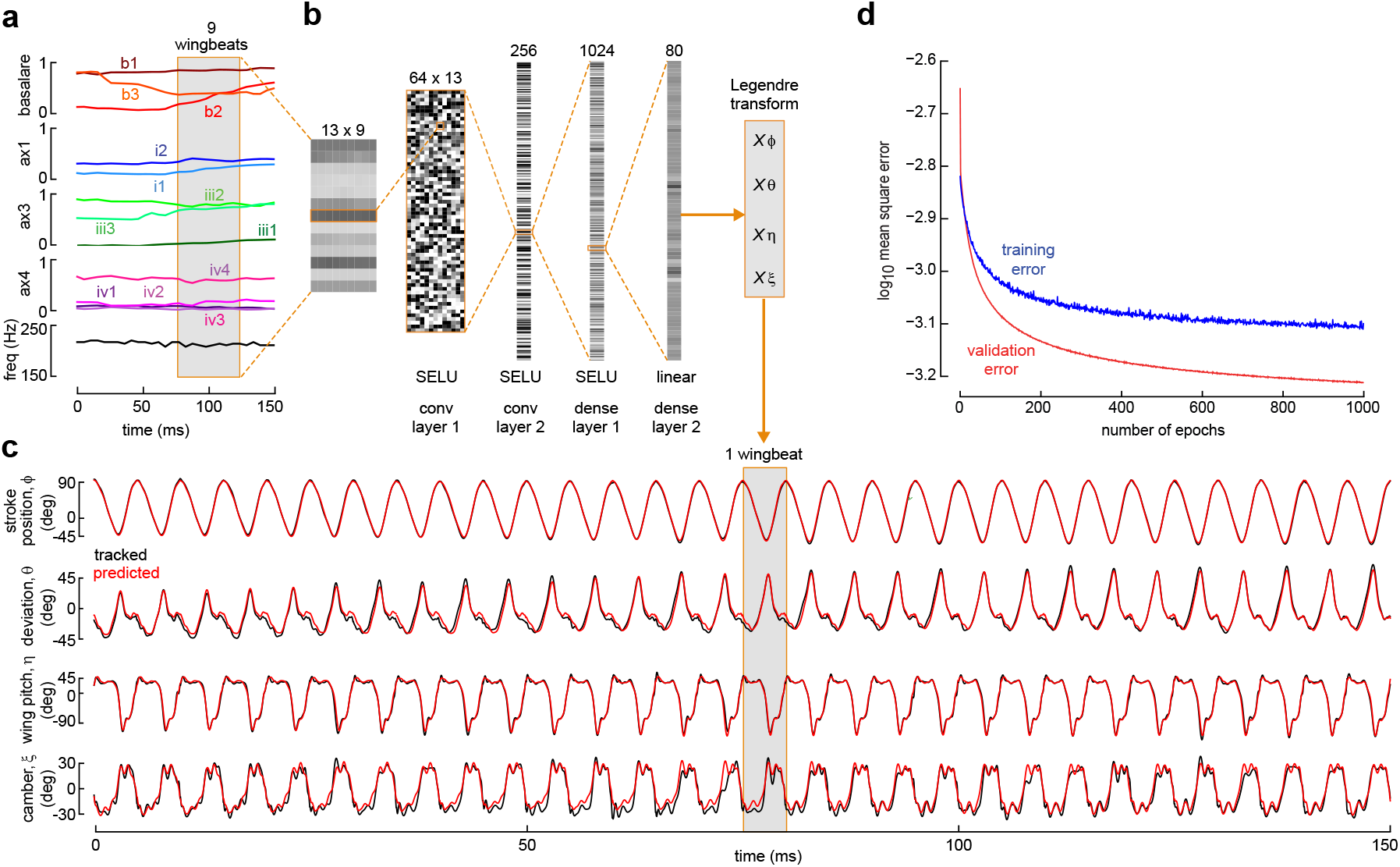
A trained CNN predicts wing motion from steering muscle activity and wingbeat frequency. **a**, Muscle activity recording displaying normalized muscle fluorescence (grouped according to sclerite) and wingbeat frequency. **b**, The trained CNN takes a 13x9 matrix as input, consisting of normalized muscle fluorescence and wingbeat frequency data over a temporal window of 9 wingbeats. Two convolutional (conv) layers with SELU activations extract features from the input matrix. A fully connected (dense) layer with 1024 neurons learns the relationships between the extracted features and an output layer of 80 neurons with linear activations. These 80 neurons correspond to 80 Legendre coefficients, encoding the four wing kinematic angles during the first wingbeat in the 9-wingbeat window. The wing kinematics can be reconstructed by multiplying the Legendre coefficients with the corresponding Legendre basis functions (Xφ, Xθ, Xη, Xξ). See Supplementary Information for more details regarding the CNN. **c**, Comparison between predicted (red) and tracked (black) traces of the four wing kinematic angles throughout the example recording. **d**, Training and validation error as a function of training epoch.

The details of the CNN that predicted wing motion from muscle activity, along with metrics used to evaluate its accuracy, are provided in Supplementary Information. Briefly, we trained the network using 85% of our dataset, reserving the rest as a test set. For each wingbeat, the 13x9 matrix of muscle activity and wingbeat frequency provided the input to the network, whereas the output consisted of a vector containing the 80 Legendre coefficients encoding the four angles of wing motion (Fig. 3a,b). Despite the fact that the network was trained using a measure of muscle activity filtered by GCaMP7f kinetics, it nevertheless predicted the salient modulations in wing motion throughout the measured sequences with remarkable accuracy (Fig. 3c, Extended Data Fig. 3). This accuracy was not guaranteed, given that prior biophysical experiments suggest that the properties of some steering muscles (most notably, b1)^20^ are regulated by small changes in motor neuron firing phase, a physiological feature that is not detectable via Ca^2+^ imaging. This success of the network might imply that such changes in firing phase are correlated with features that are detectable by our imaging method, or that excluding firing phase might be responsible for small errors in our predictions that are beyond the precision of our wing tracking. Nevertheless, the overall success of the trained network suggests that even a limited measure of steering activity provides sufficient information to reconstruct the intricate pattern of wing motion with reasonable accuracy.

A previous study found that steering muscles can be roughly categorized by their activity pattern into phasic muscles (e.g. b2, i1, iii1, iv1) that fire sporadically in bursts, and tonic muscles (e.g. b1, b3, i2, iii3, iv4) that exhibit more continuous patterns of activity^27^. We observed roughly the same pattern in our more limited dataset (Fig. 4a), which was deliberately enriched to captures sequences in which muscle activity and wing kinematics actively changed during the 1.1 ms segments during which we collected high-speed video data. To quantify the degree to which the activities of different steering muscles were linked, we performed a correlation analysis on the muscle activity, and found strong linear trends among some of the muscles (Extended Data Fig. 4, Extended Data Table 1). We did not find any significant correlation between steering muscle activity and wingbeat frequency. This was not surprising, given that changes in wingbeat frequency are known to occur more slowly than changes in the time course of the wing kinematic angles^32^, and the indirect flight muscles that have been implicated in the regulation of frequency (the power muscles^33^ and the two pleuro-sternal muscles^34^) were not recorded in this study.

**Figure 4.**
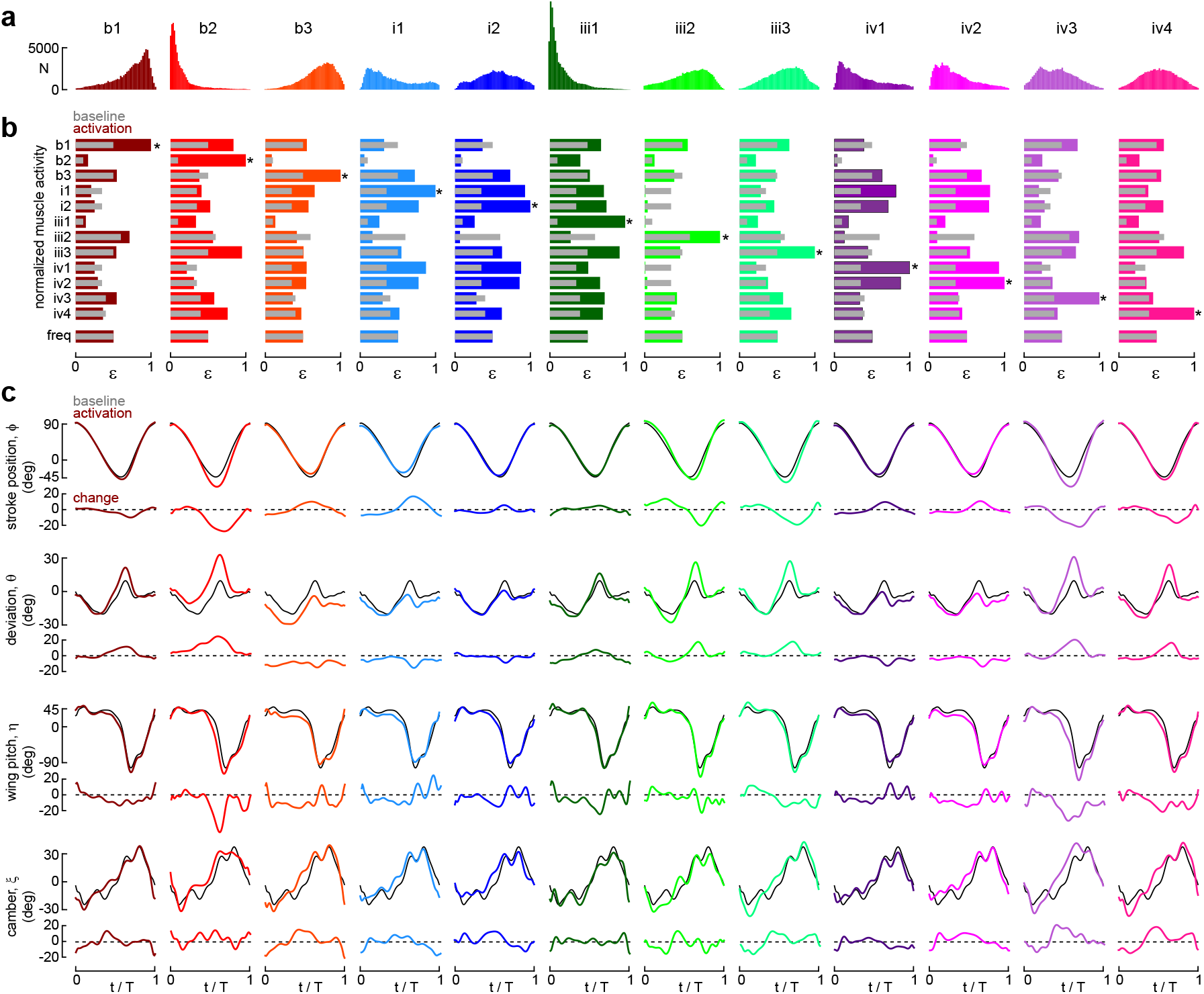
Modulation of wing motion by steering muscle activity revealed through virtual experiments. **a**, Distribution of normalized muscle activity (ϵ) for all 12 steering muscles. **b**, Muscle activity patterns generated in series of virtual experiments perform on our trained CNN model of the wing hinge. Each of the twelve set of bars beneath each muscle indicates the normalized pattern of activity across all 12 steering muscles under two conditions: 1) baseline activity when no changes in were motion were detected (inner grey bars), and 2) the activity patterns obtained by adjusting the activity of each muscle to its maximum value (outer colored bars). Note that the pattern of grey bars is identical for each muscle, because the same background pattern of activity is plotted for each case. Also note that when we changed the activity pattern of each individual muscle to ϵ=1 (marked by asterisk) we also adjusted the activity levels of all the other muscles according to the correlation patterns we observed over our entire dataset (Extended Data Fig. 4, Extended Data Table 1). This constraint helps ensure that the virtual manipulations were performed under conditions that approximated those under which the CNN was trained. Wingbeat frequency is maintained at a constant level for all patterns. **c**, CNN-predicted wing kinematics for the baseline and maximum activity patterns shown in (b) over a single wingbeat period (T). The kinematics from the baseline muscles activity patterns are shown in gray; the colored lines represent the kinematics resulting from the maximum activity patterns of the corresponding muscles; the change in kinematics induced by the simulated muscle activation is plotted as a single trace below each panel.

## Virtual manipulations using CNN model

We used our CNN model of the hinge to investigate the transformation between muscle activity and wing motion by performing a set of virtual manipulations, exploiting the network to execute experiments that would be difficult, if not impossible, to perform on actual flies. As a starting point, we defined a baseline pattern of muscle activity that was derived from sequences without any noticeable changes in muscle fluorescence and wing motion (grey data, Fig. 4b,c). From this starting point, we could then manipulate the network inputs to systematically increase the activity of each individual steering muscle, along with proportional changes in the activity of the other muscles with which it was correlated within our dataset (Extended Data Figure 4, Extended Data Table 1). The kinematic consequences of increasing the normalized activity of each steering muscle to 1, representing near-maximal output, are plotted in the columns of Fig. 4c. From these data, it is possible to identify the correlations between muscle activation and changes in wing motion, as one might do in an actual physiology experiment in which each steering muscle was recorded in isolation. The results of these virtual experiments are consistent with prior experimental data from the subset of steering muscles for which electrophysiological recordings have been possible. For example, activation of the b1 and b2 muscles is known to correlate with an increase in stroke amplitude and deviation^21,35^. Although electrophysiological recordings from b3 have not been feasible due to its small size, the antagonistic effects relative to b1 are consistent with the morphological arrangement of these two muscles. Similarly, the activity of i1 is known to correlate with a decrease in stroke amplitude and deviation, whereas the activity of the muscles of ax3 are correlated with increases in these parameters^35,36^. The consistency of our model’s predictions with these prior observations provides confidence that the CNN converged on a solution that captures the salient features of wing hinge mechanics.

## Aerodynamic effects of steering muscles

Collectively, the results of the virtual experiments conducted using our CCN hinge model indicate that the 12 steering muscles act in a coordinated manner to regulate subtle changes in wing motion. To examine the aerodynamic consequences of these changes, we used a dynamically scaled flapping robot^37^ to measure how flight forces would change with alterations in the activity of each muscle (Extended Data Fig. 5). The motion of the leading edge of the robotic wing was controlled by a system of servo motors that specified the three wing angles (*ϕ, θ, η*). Unlike earlier versions, which modeled the wing as a solid flat planform^38^, our new robotic wing consisted of 4 span-wise wedges connected by three hinge lines approximating the positions of wing veins L3, L4, and L5 (Fig. 1d, 2c). We could actuate the angles at each hinge line via servo motors attached to the base of the wing, thus specifying the fourth kinematic angle, ξ, that determines camber (Extended Data Fig. 5a). A sensor at the base of the wing measured the aerodynamic forces and torques generated by the flapping motion of each kinematic pattern. We used the Newton-Euler equations to estimate the inertial forces generated by a flapping wing, based on previously measured values of wing mass^39^ and added them to the measured aerodynamic forces, thus obtaining a time history of forces and torques generated by the wing for each wingbeat (Extended Data Fig. 5b,c). We then conducted this analysis for the 12 kinematic patterns predicted by our CNN for the maximum activity of each steering muscle (Fig. 4), creating a map linking muscle activity to flight forces (Extended Data Fig. 6).

## Simulating free flight maneuvers

Using our aerodynamic data, we could then test whether our CNN linking muscle activity to wing motion could generate free flight maneuvers that resemble those executed by real flies. To do so, we used Model Predictive Control (MPC)^40^ to determine the activity patterns within the array of 24 steering muscles (12 for each side) that generate any arbitrary flight maneuver in the shortest amount of time within a specified number of wingbeats. The details of our simulator are described in Supplementary Information (Fig. 5, Extended Data Fig. 7). Briefly, the state vector of the system, *x*, consists of 13 parameters that collectively specify the linear and angular velocity of the body in the strokeplane reference frame and the body quaternion and position in an inertial reference frame. The control vector, *u*, consists of 24 parameters representing the left and right muscle activities. Our model also incorporates body drag, the inertial and aerodynamic damping of the wings^41^, and gravitational forces. In Fig. 5a-b, we show a MPC simulation of a stereotypical free flight maneuver—a body saccade^42^—in which flies rapidly change direction in ∼100 ms^32,43^. The MPC simulation starts in forward flight mode (0.3 m s^-1^) and has to reach a goal state with a yaw rotation of 90° to the right within 10 wingbeats. Left and right steering muscle activity show relatively subtle changes during the maneuver, with the largest asymmetries in b1, b3, and iv4 activity (Fig. 5c). The saccade is achieved by a combination of roll, pitch and yaw rotations with only a slight deceleration in forward velocity (Fig. 5d). Using the time course of muscle activity, we could reconstruct the left and right wing motion patterns during the saccade using our CNN model (Fig. 5e). The asymmetries in wing motion are subtle, corresponding remarkably well to those recorded during the free flight saccades of real flies^32,43^.

**Figure 5.**
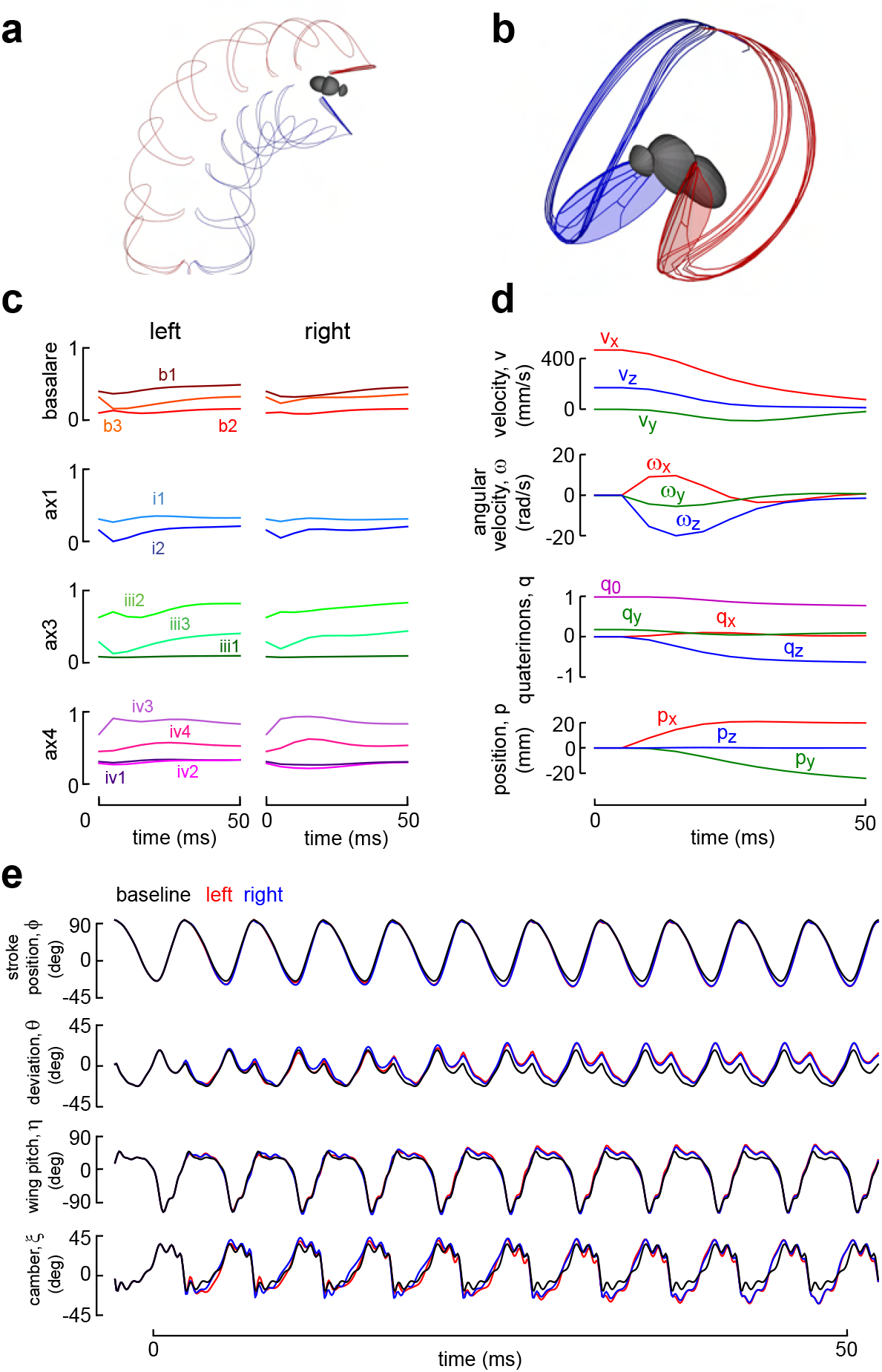
Model Predictive Control (MPC) simulation of a rightward saccade. **a**, Top view of a 30 ms simulation, showing red and blue traces for the left and right wingtips, respectively. **b**, Orthographic view of wingtip traces of the simulation in a stationary body frame. **c**, Control inputs during the saccade, representing the activity of left and right steering muscles, grouped according to sclerite. **d**, Body state during the saccade, including linear (v) and angular (ω) velocities in the SRF, as well as body quaternion (q) and position (p) relative to an inertial reference frame. **e**, CNN-predicted wing kinematic angles derived from the steering muscle activity in (c). Left wing motion is indicated in red, right wing motion in blue. The baseline wingbeat (black) is repeated throughout the maneuver for comparison. Additional MPC simulation results can be found in Extended Data Figure 7 and Supplementary Video 2.

We provide example MPC simulations of other flight maneuvers in Extended Data Fig. 7, and Supplementary Video 2. It is noteworthy that even though our CCN network of the wing hinge was trained using muscle activity data that was filtered by the time CGaMP7f kinetics, the model fly could nevertheless accomplish rapid maneuvers that closely resemble those of real flies. This result may seem counter-intuitive, but it is consistent with the known physiology of the steering muscles. While GCaMP7f dynamics are slow relative to the time course of individual muscle spikes, the more relevant physiological feature is twitch duration, which is limited by the surface area of sarcoplasmic reticulum. Even tonic muscles such as b1 that exhibit changes in dynamic stiffness due to activation phase have twitch durations lasting several wingbeats^20^. The largest saccades are associated with activation of phasic muscles b2 and i1, which act to rapidly increase and decrease wing stroke amplitude, respectively^27^. These large phasic muscles are expected to exhibit twitch durations much longer than the tonic muscles due to their specialization for maximizing rapid force production via a large cross sectional area of contractile fibers at the expense of sarcoplasmic reticulum for Ca^2+^ sequestration^44^. We suspect that our model might be less reliable for longer duration flight behaviors such as maintaining a constant heading following wing damage^22^, which require the extremely precise regulation of wing kinematics offered by the smaller, dynamically faster tonic muscles.

## Latent variable analysis

In the simulated experiments summarized in Fig. 4, we deliberately incorporated the correlation patterns among muscles (Extended Data Fig. 4), so that the results could be most accurately compared to prior work and were similar to the conditions under which we trained the network. However, the strong correlations among muscles makes it difficult to isolate their individual effects on wing motion. To gain more insight into the mechanical function of the individual wing sclerites, we constructed and trained an encoder-decoder^4^, with a network architecture that deliberately partitions the input data from the steering muscles into four nodes according to the sclerites on which they act, with an additional node assigned to wingbeat frequency (Extended Data Fig. 8). This narrow bottleneck of nodes forces the network to project muscle activity into a 5-dimensional latent-variable space, effectively performing a non-linear principal component analysis^4^. A subsequent layer of the network correlates each of these latent variables to independent changes in wing motion. After training this modified network on the dataset, we could then predict the changes in wing motion resulting from the activity of the muscle groups associated with the four wing sclerites (Fig. 6).

**Figure 6.**
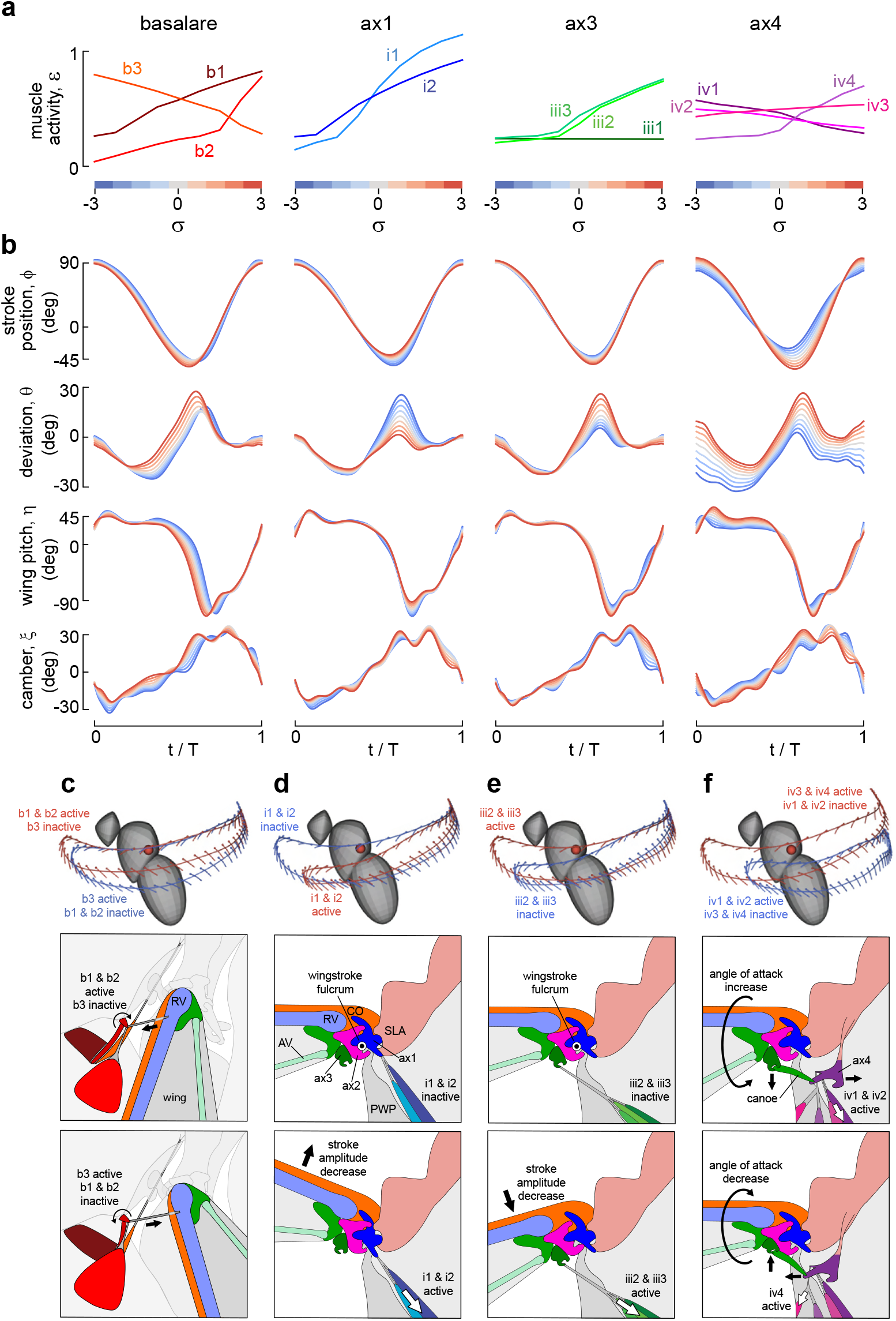
Latent variable analysis reveals the role of sclerite function in the control of wing motion. **a**, Steering muscle activity predicted by the muscle activity decoder as a function of the sclerite latent parameters (see extended Data Fig. 8). The z-scored latent variables representing each sclerite state were sampled in 9 steps, ranging from -3σ to +3σ (σ = standard deviation) and indicated by color bars. **b**, Wing kinematic angles predicted by the wing kinematics decoder, showing their dependence on sclerite state. The colors of the 9 traces correspond to the color bars in (a). **c-f**, Patterns of wing motion corresponding to extreme sclerite states (−3σ, blue; +3σ, red). Each figure plots the instantaneous wing pitch angle and wing deformation angle at regular intervals superimposed over the wingtip traces. The leading edge of the wing is indicated by a circle, and the wing’s cross-section is represented by a curved line that shows camber. **c**, Cartoon illustrating proposed mechanism of action for the basalare, drawn for a left wing from an external view. The b1 and b2 rotate the basalare clockwise, exerting tension on a ligament attached to the base of the radial vein (RV), extending the wingbeat forward and advancing wing supination during the downstroke and ventral stroke reversal. The b3 rotates the basalare counter-clockwise and has the opposite effects. **d**, Cartoon illustrating proposed action of ax1 drawn for a left wing in cross section through the winge hinge. Contraction of the i1 and i2 muscles pull ax1 ventrally, resulting in a decrease in the extent of the ventral stroke due to the inboard position of ax1 relative to ax2. **e**, Same view as in (d). illustrating action of ax3. The iii2 and iii3 muscles pull ax3 ventrally, causing an increase in the ventral stroke extent due to its outboard position relative to ax2. **f**, Same view as in (e), illustrating action of ax4. Ax4 is connected to ax3 and the Anal Vein (AV) via the long canoe sclerite. The iv1 and iv2 muscles pull ax4 inward, leading to an increased angle of attack during the downstroke via the ax4-US-ax3-AV linkage. The iv4 muscle pulls ax4 outward via its unusually thick ligament attached to the tendon of iv3, reducing the angle of attack during the downstroke.

Our analysis suggests that tension exerted on each sclerite modulates distinct aspects of wing motion (Fig. 6). For example, the b1 and b2 muscles act agonistically to increase stroke deviation and stroke amplitude during the downstroke, as well as advancing the phase of wing rotation at the start of the upstroke. The increased tension on the ligament connecting the basalare to the radial vein caused by contraction of b1 and b2 could explain these results (Fig. 6b), with the b3 muscle having an antagonistic effect. Increased downward tension on the 1^st^ axillary caused by the activation i1 and i2 decreases stroke amplitude and reduces stroke deviation at the start of the upstroke (Fig. 6d). It is noteworthy that activation of iii2 and iii3 has almost precisely the opposite effect on wing motion as those caused by activation of ax1 muscles (Fig. 6e). The positions of the ax1 and ax3 muscle insertions relative to ax2 may provide a simple explanation for these antagonistic influences on wing motion. Although the ax2 has no control muscles, this critical sclerite forms the fulcrum upon which the wing oscillates via its joint with the PWP. Whereas the muscle insertions on ax1 sits medial to this fulcrum (Fig. 6d), those on ax3 reside distally^2,9,10,45^ (Fig. 6e). Thus, whereas downward tension created by the i1 and i2 would tend to bias the mean stroke position of the wing upward (by pulling inboard of the fulcrum), tension created by the muscles of iii2 and iii3 would bias the wing downward (by pulling outboard of the fulcrum). The large iii1 muscle is rarely active during flight^27^ and is likely homologous with the ancestral retractor muscle that is used for folding the wing along the body axis^45^. When the wing is extended during the flight, however, the common tendon to which the iii2 and iii3 insert upon ax3 bend around the PWP so that the muscles exert a downward tension on the sclerite^10^.

Our latent-variable analysis suggests that the ax4 muscles exert a strong influence on wing motion (Fig 6a,b,f). In particular, the large iv1 muscle decreases the magnitude of stroke amplitude, stroke deviation, and angle-of-attack throughout the entire wingbeat, actions that are agonized by the activity of iv2, consistent with both of these muscles inserting at the same location on ax4 with a line of action that would pull the sclerite downward and inboard. In contrast, the latent variable analysis suggests that iv4 has effects opposite to that of iv1 and iv2: increasing stroke amplitude and stroke deviation and lowering angle of attack throughout the stroke, actions that are mildly agonized by iv3. These results are roughly consistent with our virtual activation experiments using the CNN model (Fig. 4). A functional stratification of the ax4 muscles into two antagonistic groups has not been observed in previous physiological studies, although these muscles have proven particularly difficult to record due to their small size and tight clustering. However, prior morphological investigations have noted that the two muscle groups insert via separate tendons that would cause different defections of ax4, which could influence wing motion via the mechanical linkage to ax3^10,46^. Our own morphological analysis (Fig. 1h, 6f, Supplementary Video 1) uncovered an intriguing feature of hinge morphology that further supports this functional stratification. In particular, the small, tonically active iv4 muscle does not insert on ax4 directly, but rather is connected laterally to the tendon of iv3 by a conspicuously broad ligament that would act to pull the sclerite outboard, antagonistic to the action of iv1 and iv2.

## Possible role of passive wing flexion

Based on our morphological and experimental results, we hypothesize that the action of the iv1 and iv2 muscles causes an inward movement of ax4 that depresses the anal vein of the wing during the downstroke via its linkage to ax3, thereby greatly increasing the angle-of-attack and reducing the lift-to-drag ratio (Fig. 6f). Whereas our the latent variable analysis predicts a large influence of ax4 muscles on stroke amplitude and deviation, the anatomical analysis of previous authors suggest that this sclerite would primarily effect the wings angle-of-attack^10^. We hypothesize that the presence of a chord-wise flexure joint at the wing root enables a passive response in the stroke and deviation angles to changes in the aerodynamic and inertial forces on the wing. Evidence for just such a flexure line has been proposed for other fly species^47^ and is evident in high speed imaging data collected during free flight in *Drosophila hydei*, which clearly show that the wing blade extends beyond the maximum angular limit of the wing base during stroke reversal, with the deformation concentrated along a narrow line (Extended Data Fig. 9a). Using confocal microscopy, we detected a strong autofluorescence signal with 405 nm excitation (Extended Data Figure 9b), consistent with the presence of the elastic protein resilin along a chord-wise line at the base of the wing blade^48^. Resilin is a rubber-like protein first discovered by Weis-Fogh in the wing hinge of locusts and flight muscle tendons of dragonflies^49^. The existence of a resilin-rich flexion joint provides a passive mechanism that could explain how alterations in the in angle-of-attack caused by activity of the ax4 muscles could generate large changes in both stroke position and deviation throughout the wingbeat, and might also play a role permitting the storage and recovery of inertial energy via elastic storage^50–52^.

## Conclusions

In this study, we used machine learning to construct both a CNN model and an encoder-decoder of the fly’s wing hinge based on the simultaneous measurement of steering muscle activity and wing kinematics. There are several reasons suggesting that our CNN model provides an accurate representation of the biomechanical processes that transform steering muscle activity into wing motion. First, when we used the network to predict the pattern of wing motion generated by the maximal activity of each muscle, the model’s output was consistent with all prior experiments for which the pattern of muscle activity has been measured directly with electrodes (Fig. 4). Second, when we embedded the CNN hinge model into a state-space simulation and used MPC to determine the pattern of muscle activity that generated an array of different canonical flight maneuvers, the results in each case resemble the known behavior of freely flying flies^32,43,53,54^ (Fig. 5). Third, when we constructed an encode-decoder in which the steering muscle activity was parsed into nodes representing the muscle groups inserting on each of the four wing sclerites (Fig. 6), the model’s predictions were consistent with known features of hinge morphology and differences in the insertion patterns of control muscles. Collectively, our results generate a suite of specific hypotheses (Fig. 6c-f) that can be tested via a combination of additional experiments and a detailed, physics-based model of the hinge. Further, with the recent availability of a connectome for the ventral nerve cord of *Drosophila*^55,56^, we expect that these results will help provide a deeper understanding of flight control circuitry that underlies the recruitment of steering muscles during flight maneuvers.

Until recently, the morphologically complex wing hinge of flies and other neopterous insects was considered a derived trait that allows the wing to fold along the body axis when not in use, and for wing motion to be driven by the contraction of indirect flight muscles that do not insert at the base of the wing. The hinge found in extant paleopterous insects (dragonflies, damselflies, and mayflies), which is actuated by direct flight muscles and does not permit folding, was considered representative of the ancestral condition^57^. However, a new phylogeny derived from a comprehensive genetic analysis of the Polyneoptera provides a compelling reinterpretation^58^, in which the unfoldable wing of the paleopterous orders is more parsimoniously interpreted as a radical simplification of the ancestral condition. According to this hypothesis, many features of the odonates—including the inability to fold their wings—are secondary specializations of their large size, perching habit, and predatory lifestyle^59–61^. Thus, despite the fact that *Drosophila* is a crown taxon, the analysis of its wing hinge could provide general insight into biomechanical innovations underling the evolution of insect flight.

## Methods

### Flies

We generated flies expressing GCaMP7f in steering muscles by crossing w[1118];+;P{y[+t7.7] w[+mC]=R22H05-Gal4}attP2 and +[HCS];P{20XUAS-IVS-GCaMP7f}attP40;+. All experiments were conducted on the 3-day old female offspring. We anesthetized flies on 4°C cold plate to immobilize them and remove the anterior two pairs of legs at the coxa. The flies were then attached to a tungsten wire at the notum using UV-curing glue (Bondic) and placed into our experimental setup after a 10-minute recovery period.

### Imaging and constructing 3D model of the wing hinge

Our reconstruction of cuticle morphology using a confocal microscopy was based on a previously published method^62^. Flies were anesthetized with acetone and briefly washed with 70% ethanol. We placed the animals in PBS with 2% paraformaldehyde and 0.1% Triton X-100 and before carefully removing their abdomen. After overnight fixation at 4°C, we isolated the thoraces and bleached the preparations for 24 hours in 20% hydrogen peroxide. The samples were then embedded in 7% agarose and cut into 0.3 mm sagittal sections using a vibratome (Leica VT1000s). The slices were incubated overnight at 37° C in PBS, 0.1% Triton X-100, and 0.2 mg ml^-1^ of trypsin to remove soft tissues. We then gradually dehydrated the preparations in a glycerol series (2% to 80%), followed by ethanol series (20% to 100%), before mounting them between two cover slips in methyl salicylate for imaging. Serial optical sections were obtained on a laser confocal microscope (Zeiss 980) at 1 µm intervals using a LD-LCI 25x, 0.8 NA objective, or at 0.3 µm intervals, using a Plan-Apochromat 40x, 0.8 NA objective. We detected the green autofluorescence characteristic of hard, sclerotized cuticle by exciting the tissue with 488 nm light. We extracted 3D meshes from the confocal stacks using a viewer plugin for *Fiji* (http://fiji.sc/) and imported the data into Blender (http://www.blender.org.). The resulting image data from the 0.3 mm sections were then processed to segment individual structures (e.g. sclerites, muscles, and apodemes) and reconstruct the continuous 3D morphology of the thorax in the vicinity of the wing hinge. The results we used to create the 2D cartoons in Fig. 1, and the animation in Supplementary Video 1.

### High-speed camera recordings

We used three synchronized high-speed cameras (SA5, Photron) to image the flies from orthogonal views. The cameras recorded continuously at a rate of 15,000 frames per second (fps) with an electronic shutter speed of 33.3 μs, telecentric lenses (0.5X PlatinumTL, Edmund Optics) and collimated infrared (850nm) backlights (M850LP1 LED and SM2P collimation lens, Thor Labs). The 3D calibration of the high-speed cameras was achieved using the Direct Linear Transformation (DLT) method^63^. The recorded images had a resolution of 256x256 pixels with 8-bit depth. To optimize data storage, we divided the camera memory buffers into 8 partitions, each accommodating one 1.1 s sequence of 16,376 frames. Once all partitions were full, the data were transferred to a hard drive. During each experiment, we also operated a low-speed tracking system called kinefly^64^, which tracked the stroke amplitude of the left and right wings so that we could implement visual closed-loop of the cylindrical array of LEDs^28^ surrounding the fly.

### Real-time calcium imaging

We employed a previously described technique^27^ to visualize muscle activity using GCaMP7f in the fly by utilizing single photon excitation through the cuticle. A blue LED (M470L3, Thor Labs) served as the excitation light source, which was directed through a 480/40 nm excitation filter (Chroma) and focused onto the fly using a 4x lens (CFI Plan Fluor, Nikon). The resulting fluorescent light passed through a 535/50 nm emission filter (Chroma) and was captured by a machine vision camera (BF3-U3-04S2M-CS, FLIR). To synchronize the imaging process, we utilized an optoelectronic wingbeat analyzer^65^ to strobe the blue excitation LED for 1 ms at the dorsal stroke reversal of each wingbeat. The fluorescence camera was strobed at half the wingbeat frequency (∼100 fps), resulting in each frame representing the sum of two consecutive illuminations. To synchronize the high-speed and fluorescence image streams, we employed a microcontroller (Teensy 3.2) that received synchronization pulses from the Photron cameras and ventral stroke reversal signals from the wingbeat analyzer, and coordinated the strobe signals to the blue excitation LED and the machine vision camera. The fluorescence images, along with a high-speed pulse count serving as a timestamp, were saved by our data acquisition software implemented in Robotic Operating System (ROS, www.ros.org) using Python scripts. An unmixing algorithm, described elsewhere^27^, was applied to extract the Ca^2+^ signal from the overlapping muscles. To initialize an experiment, a GUI was used to orient and scale the 3D muscle model to the image using an affine transformation. The resulting fluorescence image and data vector capturing the activity of the steering muscles were saved in an hdf5 file. The muscle activity vector was stored in a rolling buffer within ROS, covering approximately 30 seconds of data.

Our objective was to trigger the data sequences during rapid changes in muscle activity. To achieve this, a separate ROS node monitored the activity of a predefined muscle of interest throughout the experiment. At the start of each trial, we selected a muscle and an activity threshold for the trigger level. During the experiment, the trigger node calculated the gradient of muscle activity over 3 frames (equivalent to 6 wingbeats) and normalized this signal by dividing it by the standard deviation over a 30-s rolling buffer. Whenever the absolute value of the gradient exceeded the activity threshold, indicating a significant increase or decrease in muscle activity, the high-speed cameras were triggered in center mode, such that we saved 8,188 frames before the trigger event and 8,188 frames after. A 30-s refractory period after each trigger event ensured there was no overlap between subsequent high-speed sequences. In total, we recorded data from 82 flies, resulting in 485 high-speed videos (Extended Data Table 2). We aimed to record sequences triggered on all 12 muscles from a minimum of 5 flies each. However, the iii1 muscle exhibited sporadic activity, primarily during flight starts and stops, whereas iii2 displayed more gradual changes in activity compared to other muscles. As a result, we were only able to capture data from 1 fly for iii1 and 4 flies for iii2.

### Automated wing pose reconstruction

We developed an automated tracking system to extract body and wing pose, called Flynet, which consists of two steps: (1) a trained CNN that predicts pose vectors of the body and wing, and (2) a Particle Swarm Optimization (PSO) step that refines the CNN prediction via 3D model fitting. A GUI was created to load images, scale the 3D fly model, annotate frames, and run the automated tracking algorithm. The 3D fly model includes the head, thorax, abdomen, left wing, and right wing, each with a pose vector comprising a quaternion, *q*, and position, *p*. The CNN, built using Tensorflow (www.tensorflow.org) and Keras (keras.io) in Python 3, consists of a convolutional block to extract image features in each camera view, followed by a fully connected layer of 1024 neurons with SELU activation functions^66^ and a final layer of 37 neurons with linear activations corresponding to the pose vectors of the five 3D model components (Extended Data Fig. 2a). Further details regarding the training and operation of the automated wing tracker are provided in Supplementary Information.

We defined the Strokeplane Reference Frame (SRF) by conducting a Principal Component Analysis on the left and right wing root traces; the first, second, and third principal components of the root traces determined the y, x, and z axes of the SRF, respectively. Once the SRF was defined, we used three Tait-Bryan angles to specify the orientation of the leading edge of the wing: stroke angle φ, deviation angle θ, and wing pitch angle, η. Additionally, we defined a fourth angle, ξ, to describe chord-wise wing deformation (i.e. camber) along three flexure lines roughly corresponding to wing veins L3, L4, and L5 (Fig. 1a). We assumed equal division of the total deformation angle among these three span-wise flexure lines (Extended Data Fig. 2b).

Following the tracking process with Flynet, we applied an Extended Kalman Filter (EKF)^67^ to achieve temporal smoothing of body and wing pose. Each pose vector was independently filtered, incorporating the first and second order temporal derivatives to ensure smooth motion. The system matrix utilized the temporal derivative of quaternions and a Maclaurin series of position and wing deformation. To achieve optimal smoothing with no phase shift, we performed both a forward pass of the EKF and a backward pass using a Rauch-Tung-Striebel smoother. The Flynet GUI enables users to adjust the system covariance, allowing control over the degree of filtering for each pose parameter. To facilitate comparison between wingbeats of different durations, we normalized the wingbeat period to 1 and computed the wingbeat frequency, *f*, for each wingbeat, starting and ending with dorsal stroke reversal. To accurately quantify the complex time history of wing motion, we fit Legendre polynomials to the 4 wing kinematic angles for each wingbeat as described more fully in Supplementary Information.

### Outlier rejection, normalization, and training of the CNN

From the initial dataset of 83,056 wingbeats, we removed unrealistic flight conditions by excluding wingbeats with frequencies outside the range of 150 to 250 Hz and outliers exceeding biologically plausible angle limits for each wing kinematic angle ([-120° ≤ *ϕ* ≤ 120°], [-60° ≤ *θ* ≤ 60°], [-150° ≤ *η*≤ 150°], [-90° ≤ *ξ* ≤ 90°]). These two criteria eliminated 8.2% of the data, resulting in a final dataset of 72,219 wingbeats. Prior to training the CNN, we normalized the muscle activity dataset by dividing the GCaMP7f traces by their respective standard deviations over the entire experiment duration such that 0 corresponds to -2σ and 1 to +2σ, and normalized wingbeat frequency by mapping 150 Hz to 0 and 250 Hz to 1. These normalized values formed a 13x9 input matrix representing muscle activity and wingbeat frequency over 9 wingbeats. The output data consisted of the 80 Legendre coefficients for the left wing kinematic angles, normalized to range between -1 and 1 by dividing by π. The CNN employed a sliding window of 9 wingbeats as input and predicted the Legendre coefficients for the first wingbeat of the window. Because the muscle fluorescence was recorded at half the wingbeat frequency, we interpolated the data to predict a value for each wingbeat. We allocated the first 30 wingbeats from each high-speed video sequence for validation, while the remaining wingbeats were used for training. The training and validation datasets, consisting of 61,351 and 10,868 wingbeats respectively, were randomly shuffled. The CNN was trained for 1000 epochs using a batch size of 100, a learning rate of 10^−4^, and a learning rate decay of 10^−7^ per epoch, with the mse serving as the loss function. Gaussian noise with a standard deviation of 0.05 was added to the network input to enhance noise robustness. After shuffling the dataset following each epoch, the training and validation losses reached mse values of 10^-3.33^ and 10^-3.07^, respectively (Fig. 3d). To evaluate the accuracy, the trained CNN was used to predict wing motion for each validation sequence based on the muscle activity data. The predicted coefficient vector was multiplied by π and separated into coefficients for the four wing kinematic angles. Multiplication of the Legendre coefficient vectors by the Legendre polynomial bases recovered the four wing kinematic angles during each wingbeat. Figure 3 and Extended Data Fig. 3 illustrate examples of different sequences of wing motion predicted from muscle activity and wingbeat frequency.

### Muscle activity correlation analysis

We conducted a linear correlation analysis on the muscle activity in our dataset. For each steering muscle, we selected samples with an average gradient larger than 0.005 over the 9-wingbeat window of muscle fluorescence, a criterion based on the sharp rise and slow decay of the GCaMP7f fluorescence kernel. Scatterplots illustrating the correlations between steering muscle fluorescence and wingbeat frequency for the gradient-selected wingbeats are presented in External Data Fig. 4. By employing linear models with RANSAC fitting^68^, we determined the slopes and intercepts for all correlations. Excluding the wingbeat frequency, muscle activity can be assumed to reside on a 12-dimensional plane. To ensure that all linear correlations in muscle activity share a common origin, we defined a baseline muscle activity pattern. This involved identifying sequences in the dataset with no significant changes in muscle activity or wing motion and setting the wingbeat frequency to 200 Hz. In Extended Data Fig. 5, the intercepts of the linear model were adjusted such that the baseline muscle activity pattern aligns with the 12-dimensional plane (Extended Data Table 1).

### Virtual experiments

To conduct virtual experiments on the wing hinge (Fig. 4), we utilized the measured muscle activity correlation (Extended Data Fig. 4, Extended Data Table 1) in conjunction with the trained CNN. In order to obtain realistic inputs to the CNN, we used a linear model to find steering muscle activity patterns that were within the manifold of the dataset that was used to train the CNN. First, we determined the baseline wing kinematics through the CNN’s prediction of the wing motion corresponding to the baseline pattern of muscle activity repeated over 9 wingbeats. To find the maximum muscle activity patterns for each muscle, we traversed the 12-dimensional surface from the baseline muscle activity pattern to a point where the specific muscle reached an activity value of 1.

### Robotic wing experiments

The aerodynamic forces generated by the baseline and maximum muscle activity patterns predicted by the CNN were assessed using a dynamically scaled robotic wing consisting of a stepper motor (IMS M-2231-3) controlling *ϕ* through two gears with a 1:3 ratio, and two servo motors (HiTec D951TW) controlling *θ* and *η*. A 6-degree-of-freedom sensor (ATI Nano 17) captured forces and torques along three orthogonal axes at the wing’s base. A microcontroller (Teensy 3.2), operating at 100 Hz, updated stepper and servo positions and collected sensor data (Extended Data Fig. 5a). The wing itself consisted of four acrylic panels and three micro-servos (HiTec HS-7115TH) at the base, collectively controlling the fourth wing kinematic angle, ξ. Each trial involved repeating the programmed motion pattern for seven wingbeats. The commands for the first and last wingbeats of each sequence were multiplied by the first and second quarter, respectively, of a sin^2^(t/T) function, ensuring that the wing began and ended at the home position (*ϕ, θ, η, ξ* = 0). To reach steady-state conditions for wake effects, the measurements from the first three wingbeats were discarded^69^, and we calculated median force and torque values over wingbeats 4, 5, and 6. In a separate experiment, gravitational and buoyancy forces acting on the wing were measured by playing back the wing motion at a 0.2X speed, enabling subtraction of the gravity measurement from the corresponding 1X speed experiment to isolate aerodynamic forces and torques. As described in Supplementary Information, the values for total forces and torques that we report (*F*_total_, *T*_total_) are the sum of aerodynamic components measured using the dynamically scaled wing and inertial components calculated using the Newton-Euler equations.

### Model Predictive Control simulations

To simulate free flight maneuvers, we integrated a non-linear, discrete-time, state-space model of a flying fly, described in the Supplementary Information, into a Model Predictive Control^70^ (MPC) loop using the do-mpc Python package^40^ (Extended Data Fig. 7, Supplementary Video 2). Each MPC simulation involved defining a start state (*x*_init_), a goal state (*x*_goal_), and a time period for the maneuver execution. The optimization process computed the trajectory that minimizes the mse between the current state and the goal state, aiming to reach the goal state in the shortest possible time. The MPC controller utilized dynamic programming to optimize the state-space trajectory over a finite horizon, considering the objective cost function and non-linear constraints. In our simulations, the finite horizon was set to 10 wingbeats, ensuring continuous trajectory computation toward the goal state. The control inputs were the activity patterns of the left and right steering muscles. We restricted the muscle activity patterns based on the correlation analysis shown in Extended Data Fig. 4. Specifically, we limited the steering muscle activity to lie within 10% of the normalized 12D muscle activity plane for both wings. The muscle activity of each muscle was also bounded between -0.2 and 1.2. We assigned a penalization weight of 1 for each muscle was set to 1, with the exception of muscles b2 and iii1, which were assigned penalization weights of 2 and 10, respectively. These higher weights were chosen because muscles b2 and iii1 are so seldom active^27^. In a similar fashion, the importance of the different components of the objective function was be adjusted by setting weights. For instance, for symmetrical maneuvers (e.g. forward acceleration, ascent, descent), we assigned higher weights to state variables that break symmetry.

### Latent variable analysis

As described more fully in the Supplementary Information, we employed an encode-decoder architecture for latent variable analysis^4^, in which the latent variables predicted both muscle activity and wing motion (Extended Data Figure 9a). The encoder split the input data into five streams, each processed by a separate CNN, to project onto an individual latent variable. The muscle activity decoder reconstructed the input matrix using a fully-connected dense layer with hyperbolic tangent (tanh) activation and a deconvolutional layer with SELU activation. To focus on the effects of sclerite state on wing kinematics, we introduced a back-propagation stop between the latent space and the muscle activity decoder. Instead of relying on the muscle activity reconstruction error, we used the wing kinematics decoder’s reconstruction mse to update the encoder weights. The wing kinematics decoder utilized a dense layer of 1024 neurons with SELU activation to predict the 80 Legendre coefficients representing wing motion during a wingbeat. After training the latent network for 1000 epochs with mse as the error function, the training error for wing kinematic reconstruction was mse = 6.0 · 10^−4^, and the validation error was mse = 7.9 · 10^−4^ (Extended Data Fig. 2d). The changes in wing motion predicted by variation in the latent variables associated with each wing sclerite (Fig. 6b) were roughly consistent with the results of our virtual muscle activation experiments (Fig. 4c). Besides sclerite functionality, the latent variable analysis also predicts the effect of wingbeat frequency on wing motion. Our analysis predicted a decrease in stroke amplitude and downstroke-to-upstroke ratio for increasing frequency, consistent with a study on the power requirements of forward flight^39^. In Extended Data Fig. 8d, we present four aerodynamic parameters: absolute angle of attack, wingtip speed, and non-dimensional lift and drag in the SRF, where the quasi-steady model was used to compute lift and drag forces. The computed lift and drag values were lower than the measurements from the dynamically scaled model wing, a discrepancy that might be due to the omission of wing deformation in the quasi-steady calculations.

## Acknowledgements

The authors wish to thank several individuals whose efforts contributed to the work presented in this paper. Will Dickson provided extensive expertise in instrumentation, programming, data analysis, and formatting all the data and code for public repositories. Thad Lindsay assisted in the design of the epifluorescent microscope and data acquisition software used for muscle imaging. Anne Erickson gave helpful comments on the manuscript and Supplementary Information. Ainul Huda assisted in the construction of genetic lines. Jaison Omoto made confocal images of wings. John Tuthill and Tony Azevedo offered a tomographic dataset of the *Drosophila* wing hinge that was collected at the European Synchrotron Radiation Facility in Grenoble, France. Sam Whitehead analyzed this tomography data to provide a preliminary reconstruction of the hinge sclerites, and offered critical feedback on the manuscript text and data presentation. Benjamin Fabian and Rolf Georg Beutel provided micro-CT data from their publication on the morphology of the adult fly body. The research reported in this publication was supported by the National Institute of Neurological Disorders and Stroke of the NIH (U19NS104655).

## Author Contributions

Johan M. Melis collected all the data presented in the manuscript and developed the software for data analysis. Johan M. Melis and Michael H. Dickinson collaborated on planning the experiments, preparing figures, and writing the manuscript. Igor Siwanowicz collected the high-resolution morphological images of the *Drosophila* thorax and created Supplementary Video 1.

## Competing Interest

The authors have no competing interests to report.

## Financial Interests

The authors have no financial interests to report.

Correspondence and requests for materials should be addressed to Michael H. Dickinson at.

## Extended Data Figure Legends

**Extended Data Figure 1.**
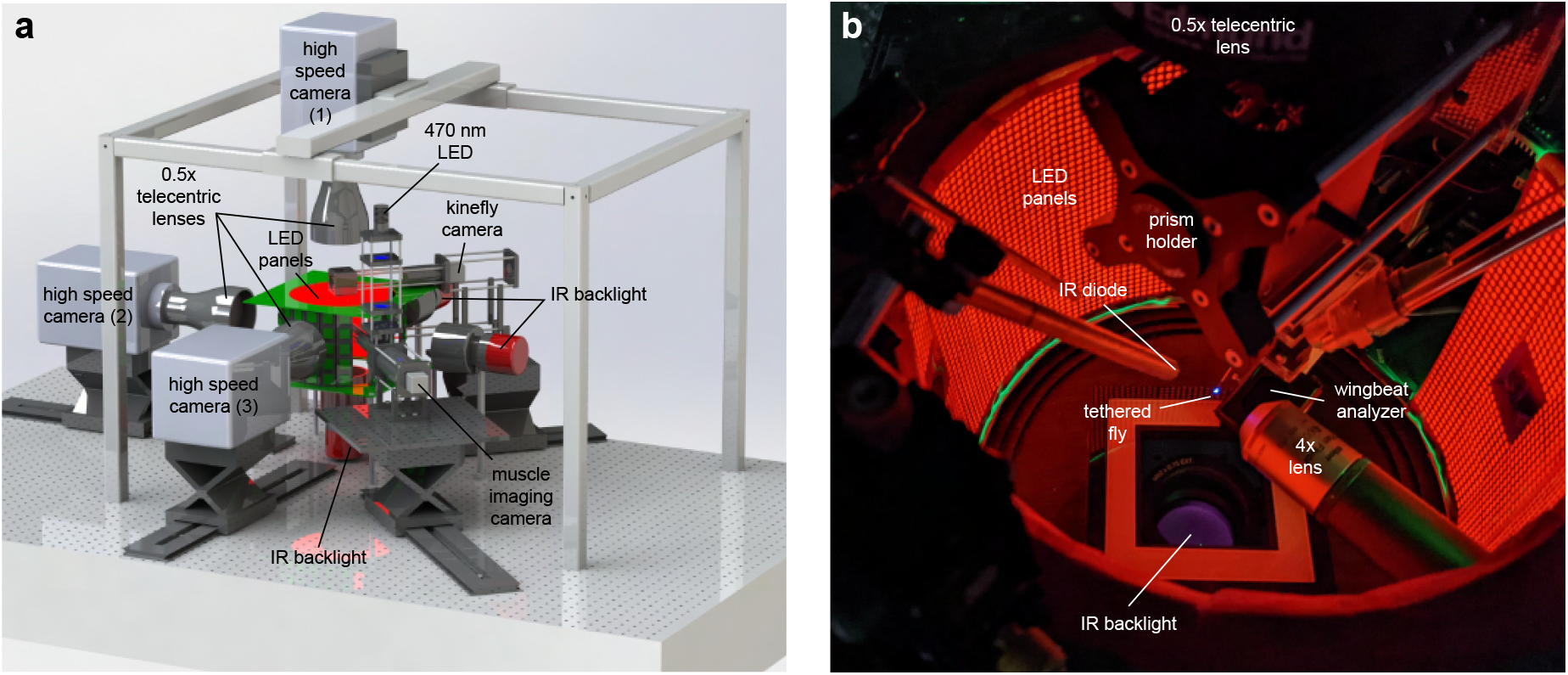
Automated setup for simultaneous recording of muscle fluorescence and wing motion. **a**, High-speed cameras, equipped with 0.5X telecentric lenses and collimated IR back-lighting capture synchronized frames of the fly from three orthogonal angles at a rate of 15,000 frames per second. An epi-fluorescence microscope with a muscle imaging camera records GCaMP7f fluorescence in the left steering muscles at approximately 100 frames per second, utilizing a strobing mechanism triggered every other wingbeat. A blue LED provides a brief, 1 ms illumination of the fly’s thorax during dorsal stroke reversal. A camera operating at 30 fps captures a top view of the fly for the kinefly wing tracker. **b**, Image of the flight arena featuring the components of the setup: LED panorama, IR diode and wingbeat analyzer for triggering the muscle camera and blue LED, prism for splitting the top view between the high-speed camera and kinefly camera, IR backlight, 4X lens of the epi-fluorescence microscope, and a tethered fly illuminated by the blue LED.

**Extended Data Figure 2.**
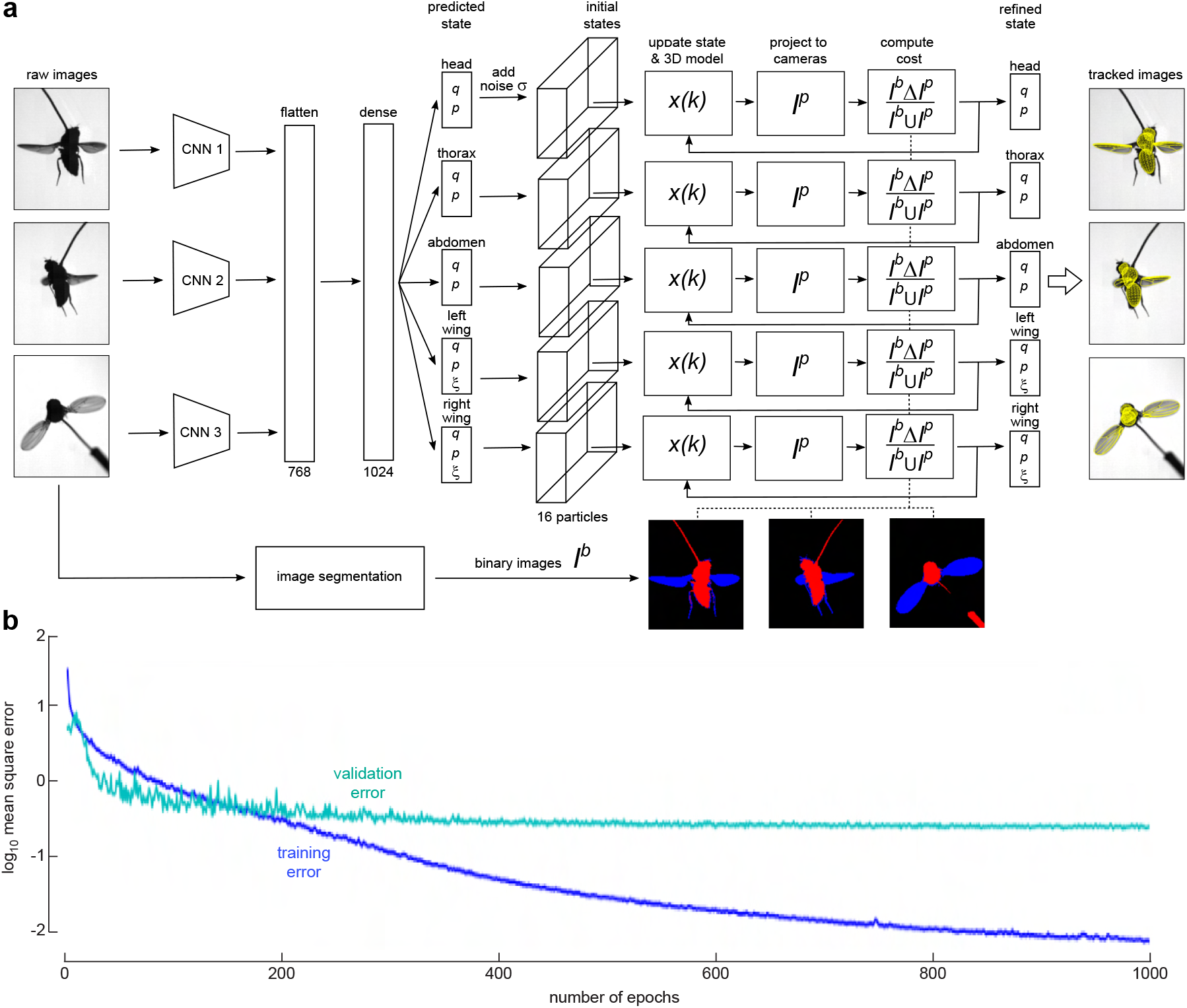
Flynet workflow and definitions of wing kinematic angles. **a**, The Flynet algorithm takes three synchronized frames as input. Each frame undergoes CNN processing, resulting in a 256-element feature vector extracted from the image. These three feature vectors are concatenated and analyzed by a fully connected (dense) layer with Scaled Exponential Linear Unit (SELU) activation, consisting of 1024 neurons. The output of the neural network is the predicted state (37 elements) of the five model components represented by a quaternion (q), translation vector (p), and wing deformation angle (ξ). Subsequently, the state vector is refined using 3D model fitting and particle swarm optimization (PSO). Normally distributed noise is added to the predicted state, forming the initial state for 16 particles. During the 3D model fitting, the particles traverse the state-space, maximizing the overlap between binary body and wing masks of the segmented frames (*I*^b^) and the binary masks of the 3D model projected onto the camera views (*I*^p^). The cost function (*I*^b^ Δ*I*^p^)/(*I*^b^∪*I*^p^) is evaluated iteratively for a randomly selected 3D model component. The PSO algorithm tracks the personal best cost encountered by each particle and the overall lowest cost (global best). After 300 iterations, the refined state is determined by selecting the global best for each 3D model component. See Supplementary Information for more details. **b**, Training and validation error of the Flynet CNN as a function of training epoch.

**Extended Data Figure 3.**
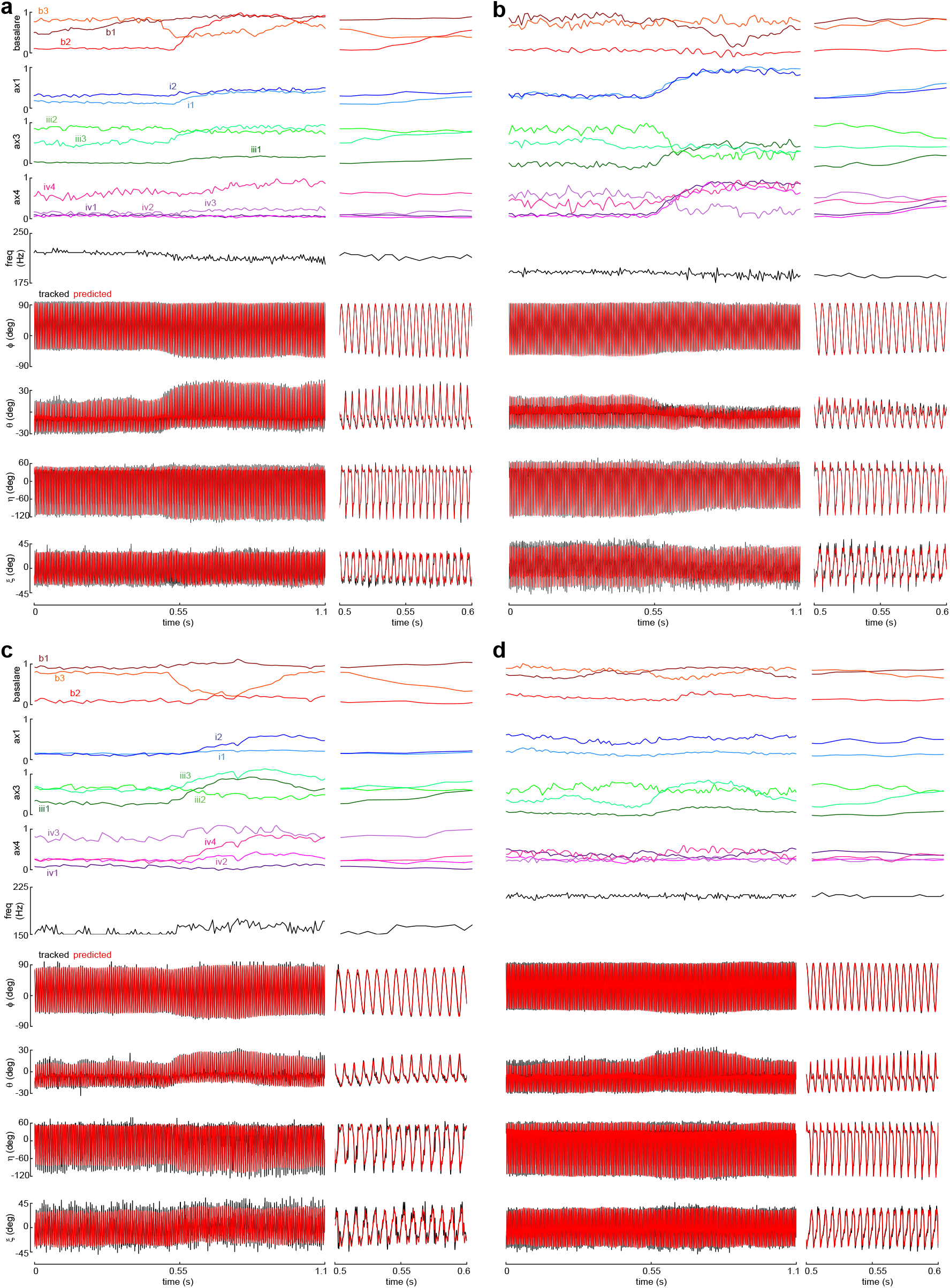
CNN-predicted wing motion for example flight sequences. **a**, The top five traces show activity of the steering muscles in the four sclerite groups as well as wingbeat frequency during a full, 1.1 second recording. The bottom four traces indicate comparison between the tracked (black) and CNN-predicted (red) wing kinematic angles throughout the sequence. Expanded plots of a 100-ms sequence (0.5 to 0.6 seconds) are plotted on the right. **b, c, d**. Same as but for a different flight sequences.

**Extended Data Figure 4.**
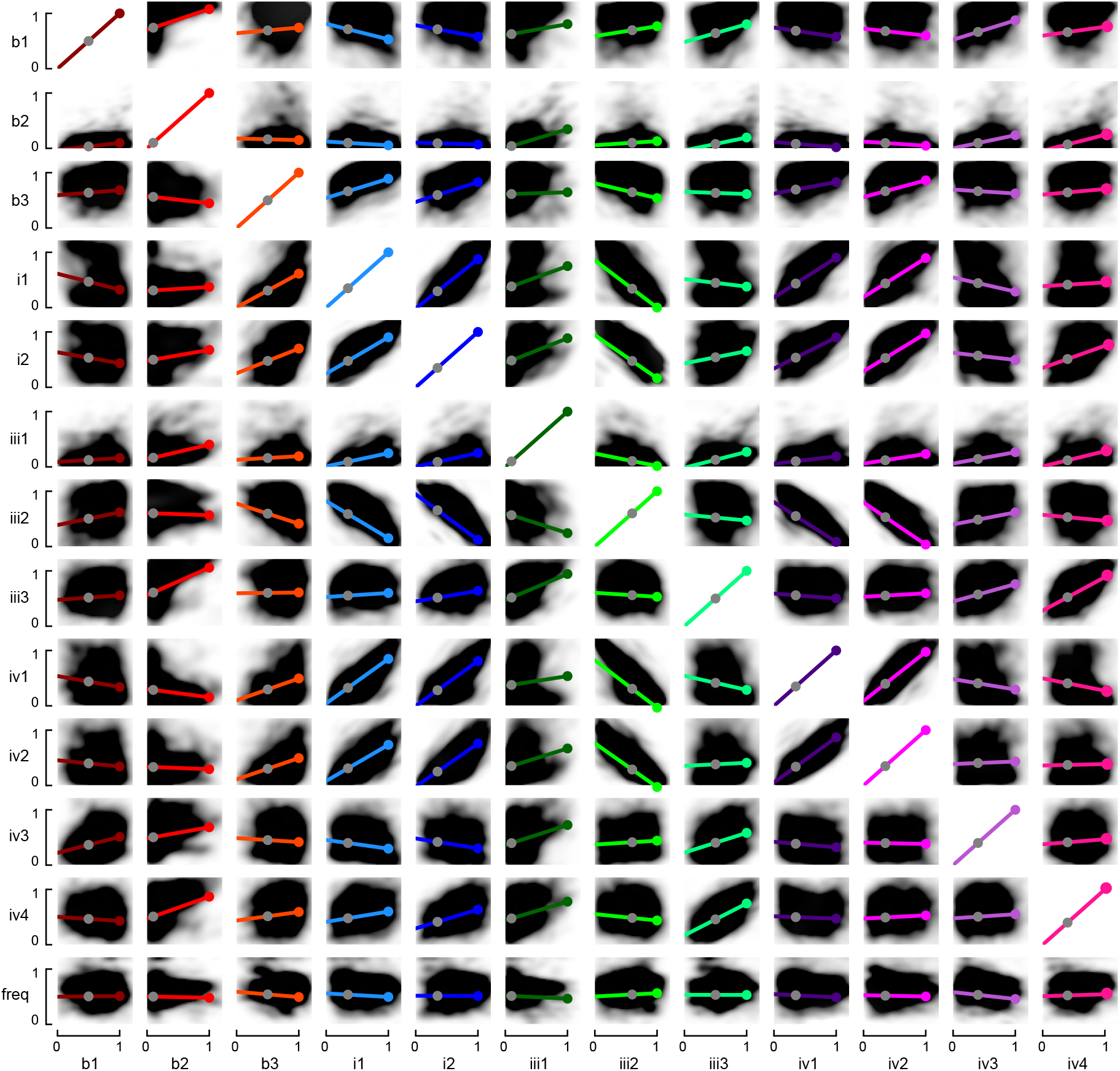
Correlation analysis of steering muscle fluorescence and wingbeat frequency. Linear models (colored lines) fitted to wingbeats in the entire dataset of 72,219 wingbeats from 82 flies. Gray dots represent the normalized baseline muscle activity level, while colored dots represent the normalized maximum muscle activity level. The correlation coefficients associated with these plots are provided in Extended Data Table 1. For more detail on regression methods, see Supplementary Information.

**Extended Data Figure 5.**
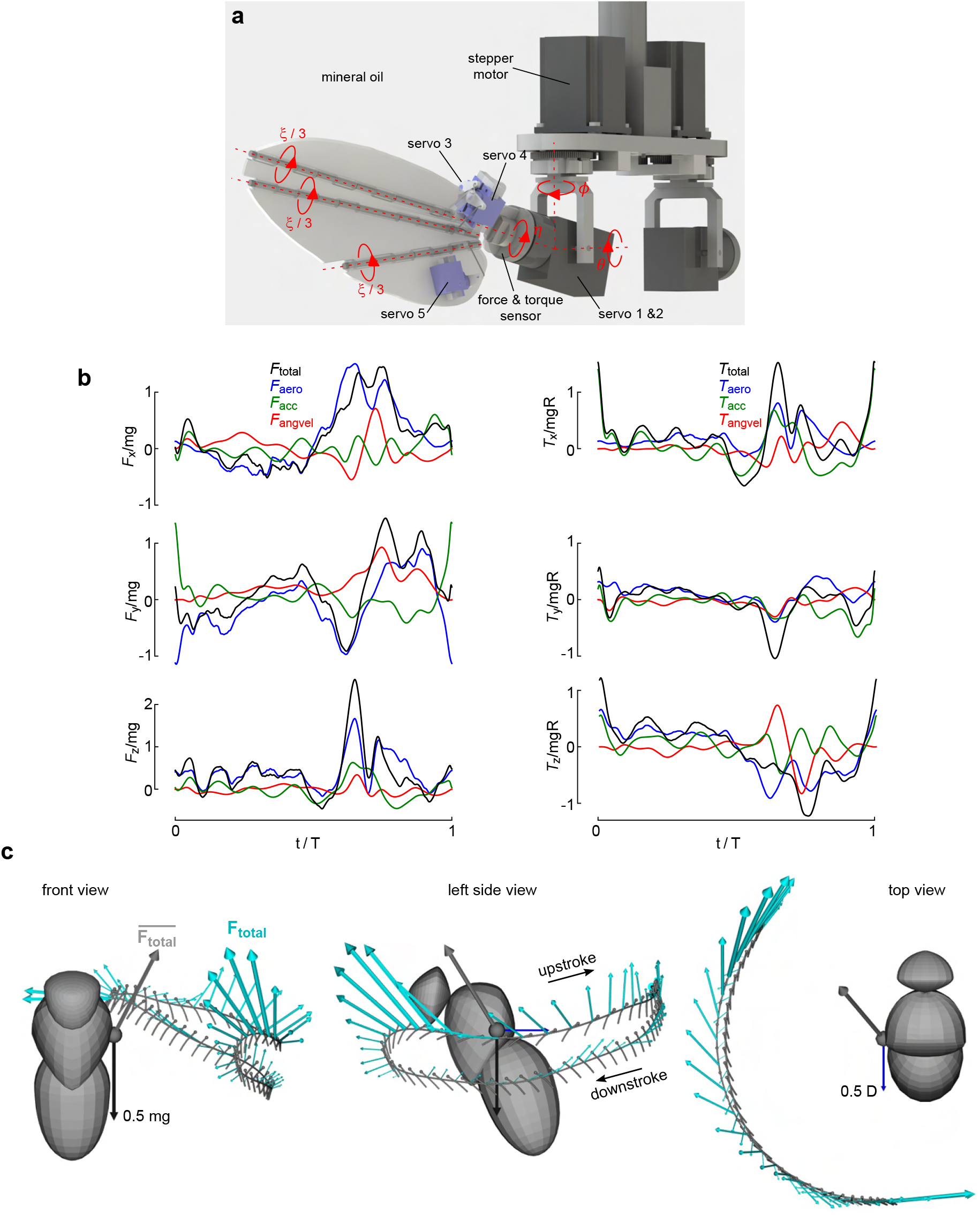
Aerodynamic force measurements and inertial force calculations. **a**, Dynamically scaled flapping fly wing model immersed in mineral oil. **b**, Non-dimensional forces and torques in the SRF for the baseline wingbeat. The four traces in each panel correspond to the total (black: *F*_total_, *T*_total_), aerodynamic (blue: *F*_aero_, *T*_aero_), inertial components due to acceleration (green: *F*_acc_, *T*_acc_), and inertial components due to angular velocity (red: *F*_angvel_; *T*_vel_). See Supplementary Information for more details. **c**, Representation of total forces during the baseline wingbeat, viewed from the front, left, and top. Gray trace represents the wing trajectory; cyan arrows represent instantaneous total force on the wing. At the wing joint, three arrows depict the total mean force, half the body weight, and half the estimated body drag.

**Extended Data Figure 6.**
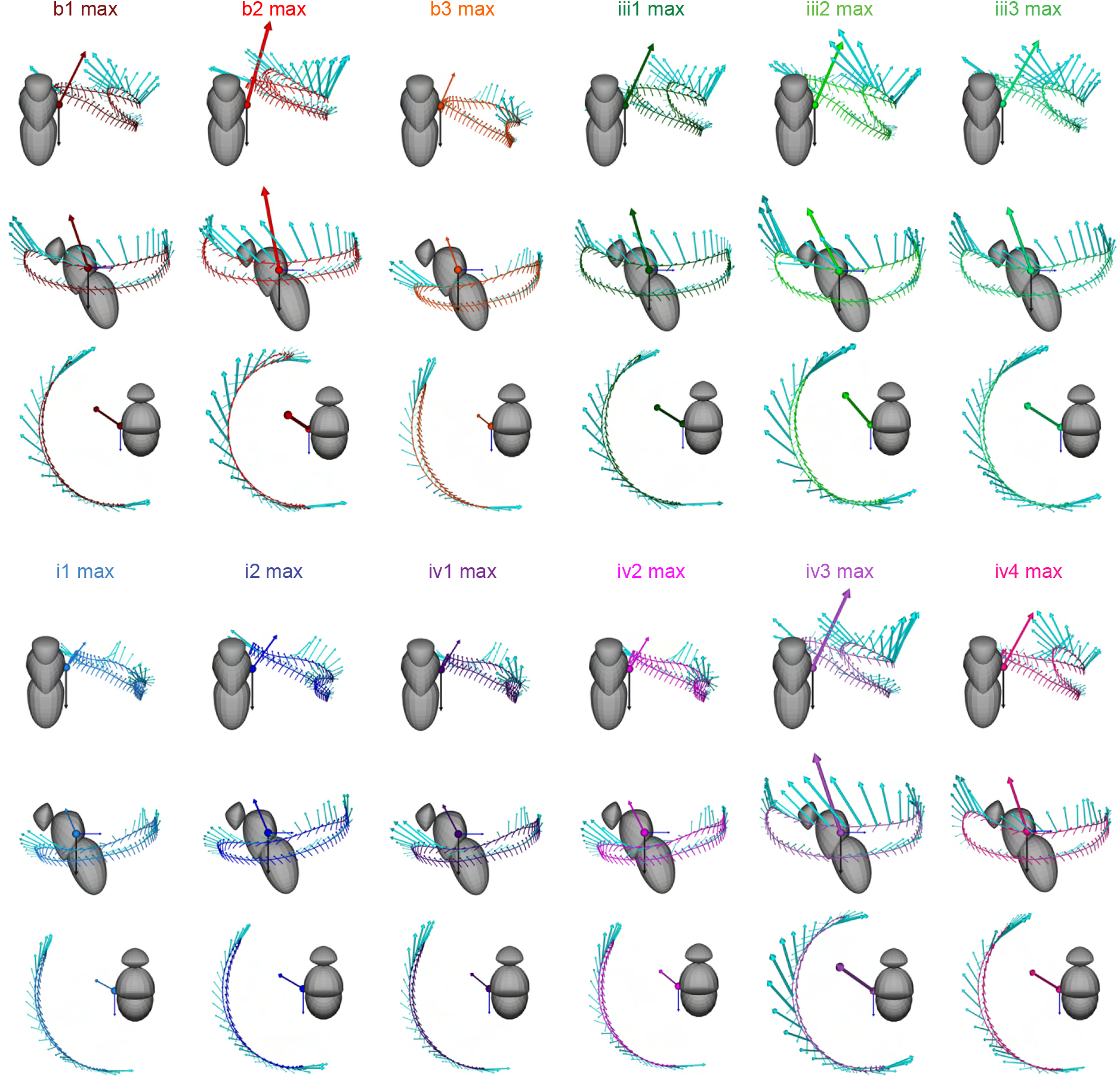
Aerodynamic and inertial forces for maximum muscle activity wingbeats. Figures depict the CNN-predicted wing motion for maximum muscle activity patterns, viewed from the front, left, and top. Instantaneous vectors depicting the sum of aerodynamic and inertial forces are shown in cyan. The wingbeat-averaged force vector is indicated by the color corresponding to the specific steering muscle set to maximum activity. Note that the scaling for the wingbeat-averaged forces differs from that for the instantaneous forces. The black gravitational force and blue body drag force are plotted as in Extended Data Fig. 6c.

**Extended Data Figure 7.**
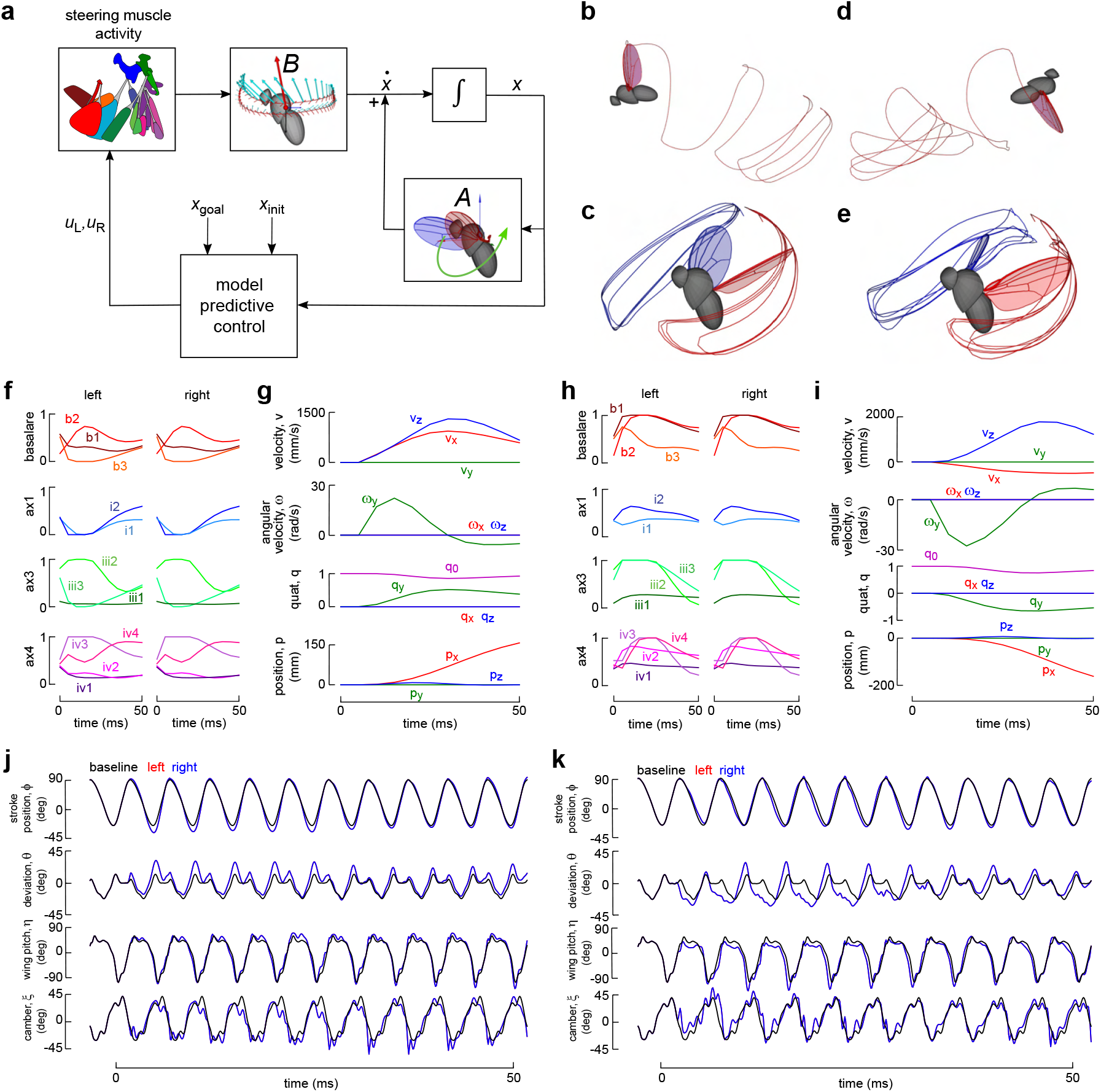
Simulation of free flight maneuvers using the state-space system and Model Predictive Control. **a**, Schematic of the state-space system and MPC loop, including system matrix (*AA*), control matrix (*B*), the state vector (*x*), temporal derivative 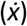 left and right steering muscle activity (*u*_L_, *u*_R_), initial state (*x*_init_) and goal state (*x*_goal_). **b**, Forward flight simulation with wingtip traces in red and blue. **c**, Wing motion during forward flight simulation plotted in stationary body frame. **d**, Backward flight simulation. **e**, Wing motion during backward flight simulation plotted in stationary body frame. **f**, Left and right steering muscle activity during the forward flight maneuver. **g**, State vector during forward flight maneuver. **h**, Steering muscle activity for the backward flight maneuver. **i**, State vector for the backward flight maneuver. **j**, CNN-predicted left (red) and right (blue) wing kinematics for the forward flight maneuver. Note that the left wing kinematics are displayed underneath the right kinematics, and a baseline wingbeat is shown to emphasize the relative changes in wing motion. **k**, CNN-predicted wing motion for the backward flight maneuver.

**Extended Data Figure 8.**
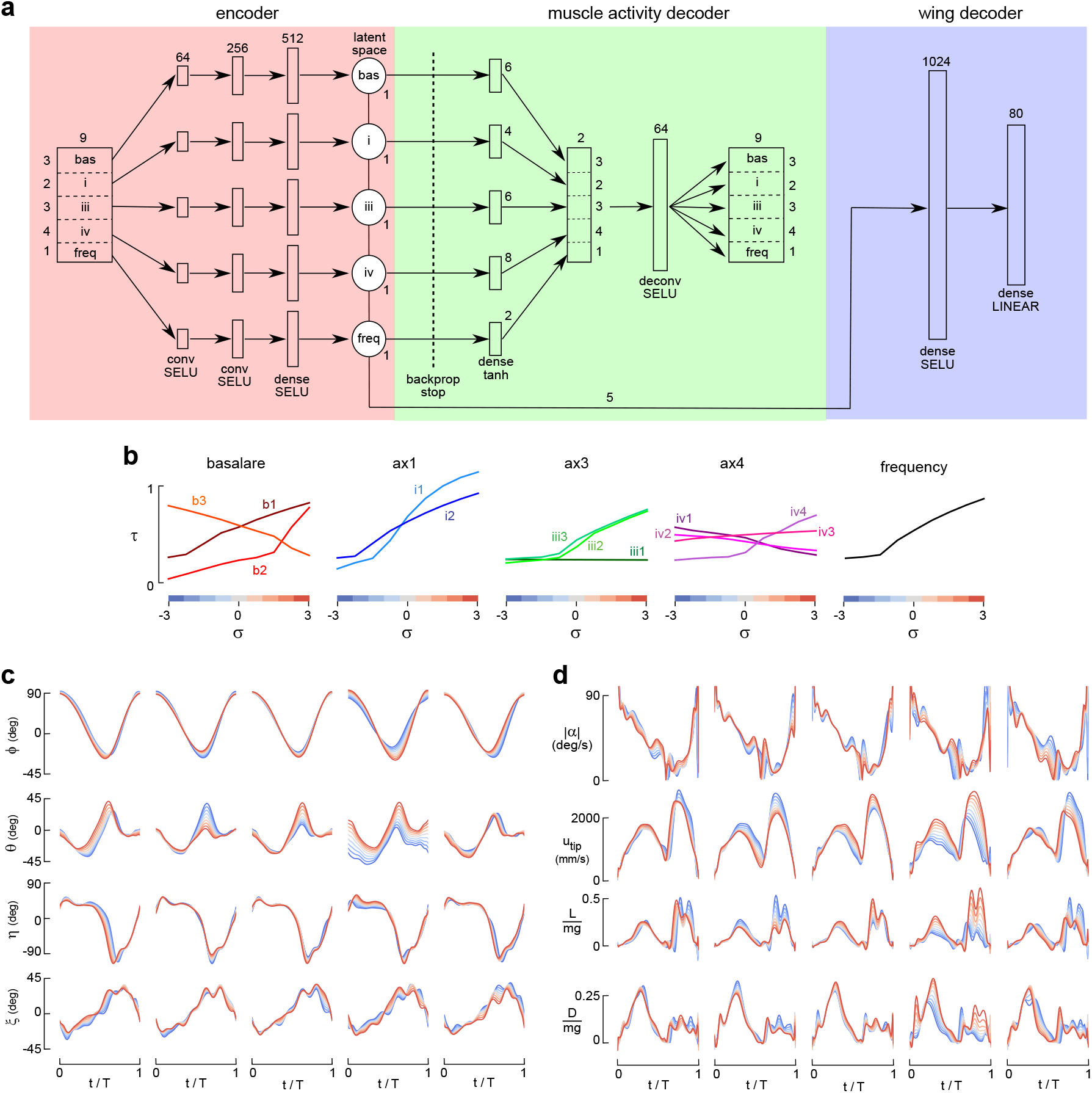
Latent variable analysis reveals sclerite function using an encoder-decoder. **a**, The network architecture consists of an encoder (red), muscle activity decoder (green), and wing kinematics decoder (blue). The encoder splits the input data into five streams corresponding to different muscle groups and frequency. Feature extraction is performed using convolutional and fully connected layers with SELU activation. Each stream is projected onto a single latent variable. In the muscle activity decoder, the latent variables are transformed back into the input data. A backpropagation stop prevents weight adjustments in the encoder based on the muscle activity reconstruction. The wing kinematics decoder predicts the Legendre coefficients of wing motion using the latent variables. See Supplementary Information for more details. **b**, Predicted muscle activity and wingbeat frequency as a function of each latent parameter varied within the range of -3σ to +3σ. Color bar indicates the latent variable value in panels (c) and (e). **c**, Predicted wing motion by the wing kinematics decoder for the five latent parameters. **d**, Absolute angle of attack (|α|), wingtip velocity (*u*_tip_) in mm s^-1^, non-dimensional lift (L mg^-1^), and non-dimensional drag (D mg^-1^). The non-dimensional lift and drag were computed using a quasi-steady model.

**Extended Data Figure 9.**
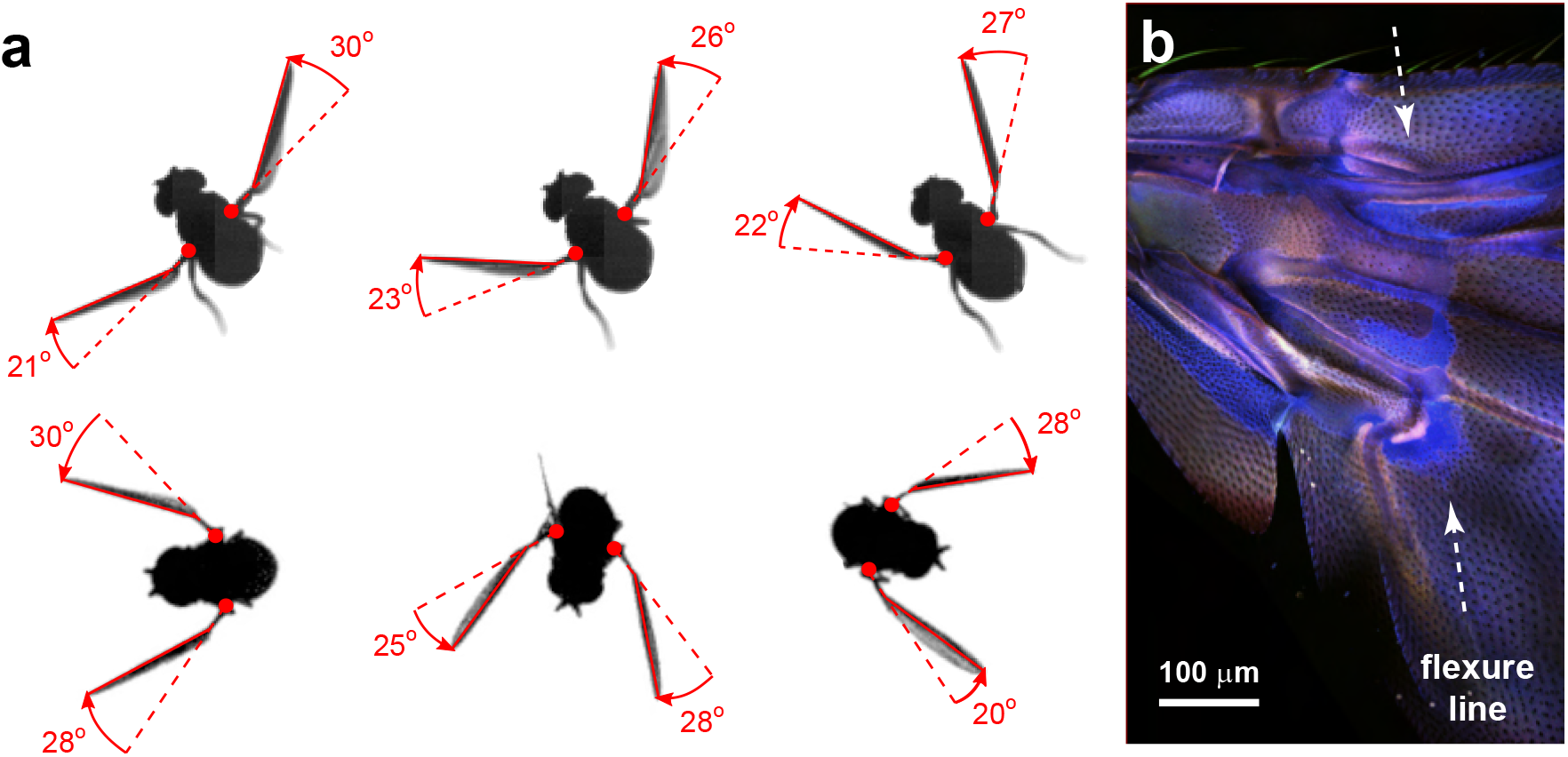
Flexible wing root facilitates elastic storage during wingbeat and allows wing to passively respond to changes in lift and drag throughout stroke. **a**, Top view of ventral stroke reversal in tethered and free flight. Red circles mark the estimated position of the wing hinge, dotted lines indicate the expected position of the wing if a chord-wise flexure line was not present. Images are reproduced from previously publish data^32^. **b**, Composite confocal image of the wing base, indicating a bright blue band of auto-fluorescence consistent with the presence of resilin and existence of a chord-wise flexure line (dashed arrows).

## Supplementary Video Legends

**Supplementary Video 1:** The left side of a Drosophila thorax, annotated to illustrate the arrangement of wing sclerites and associated musculature; color scheme is consistent with Figure 1.

**Supplementary Video 2:** Animations of simulated flight maneuvers shown in world and body frames (forward, ascent, descent, sideways, left saccade, and right saccade) generated using the CNN model of the wing hinge and state-space model operating with a Model Predictive Control loop (see Fig. 5, Extended Data Fig. 7).

## Supplementary Information

### Initial prediction of wing pose with using a convolutional neural network (Flynet)

Our pose estimation strategy required constructing an anatomically accurate 3D model of a fly that we would fit, frame-by-frame, to the data in our three high speed video sequences. We based our 3D model from images of a tethered flying fly taken at different angles, from which we extracted aspect ratios and curvatures of the head, thorax, and abdomen. These 2D contours were converted into 3D surfaces via Non-Uniform Rational B-splines (NURBS)[1] Like B-splines, NURBS surfaces may be manipulated by changing the locations of control points. By collapsing control points at the edges of NURBS surfaces onto each other, we created spheres for the head, thorax, and abdomen and then manipulated the control points to reshape the objects according to the aspect ratios and curvatures derived from the fly images.

Observations from high-speed videos show that the wings exhibit chord-wise deformations during flight that are concentrated at the L3, L4, and L5 veins (Fig. 1d). Accordingly, our model wing consists of four rigid panels connected by three hinge lines. To reduce the number of parameters representing wing deformation, we assumed that the deformation angle along each hinge line was equal (Fig. 2c). We thus modeled the total deformation angle, ξ, as the sum of the local deformation angle of each hinge line (ξ = ξ / 3 + ξ / 3 + ξ / 3), an assumption that was supported by manual inspection of the raw images.

Each wing pose vector consisted of 8 values: 4 parameters for a quaternion describing the orientation of the leading edge panel of the wing relative to the world reference frame, 3 parameters for a translation vector describing the distance between the wing root and the origin of the world reference frame, and the wing deformation parameter ξ. For the head, thorax, and abdomen pose vectors we used 7 parameters: 4 for the quaternion and 3 for the translation vector. Our total fly pose vector thus required 37 parameters to account for the head (7), thorax (7), abdomen(7), left wing (8), and right wing (8). Besides this pose vector, we defined a vector that contained 5 parameters for scaling the body and wing components independently.

We developed software, called Flynet, to implement the entire wing pose estimation process, including the manual annotation required for creating a training set. A module in the Flynet GUI constructed in Python with the PyQt and PyQtGraph packages allows the user to load the 3D fly model and scale, rotate, translate, and deform all the components. The GUI uses the DLT-calibration[2] of the high-speed cameras to project a wireframe of the fly model onto the three camera views. By adjusting the pose and scaling vectors of each component until it matches the images in each view, an accurate label can be created for each frame triplet. Using this GUI, we collected a manually annotated a final dataset of 2905 frame triplets and associated pose vectors. Neural networks work best with data centered around 0 with a standard deviation of 1. Thus, the 8-bit pixels of the high speed images were rescaled by dividing by 255. To reduce computation time, the normalized frames were cropped into 224 × 224 pixel images centered around the thorax of the fly.

The CNN was implemented in Python using the Tensorflow library with Graphics Processing Unit (GPU) support. The CNN architecture started with three convolutional blocks, one for each camera view. A graphical representation of the complete Flynet code (created using the the model plotting utility in Keras 3.0) is shown in Appendices A and B; the full code is available as described in the Data Availability Section of the main manuscript.

We randomly split our set of manually annotated data into groups of 95% and 5% for for training an validation, respectively. The network was trained for 1000 epochs with a batch size of 50. After some testing, we chose a dropout rate of 0.1 as it gave the lowest validation error. Lower dropout rates reduced the training error but increased validation error; higher dropout rates increased the training error and therefore also the obtainable validation error. Mean squared error (MSE) served as the loss function. We used a learning rate of 1.0 · 10^−4^ and a decay of 1.0 · 10^−7^. The decay value was subtracted from the learning rate after each epoch, resulting in a linear decrease of the learning rate during training. Using a Nvidia Geforce GTX 1080 graphics card with 8 GB of memory, it took approximately 24 hours to train Flynet. During the training phase, the MSE for the independent components of the pose vector decayed up to 400 epochs and remained constant afterwards (Extended Data Fig. 2b). Stabilization of the validation error indicates that the network’s performance did not improve with more epochs, even though the training error continued to decline. The weights of the trained network were saved in an hdf5 file and loaded in the Flynet GUI for the pose prediction.

### Refining pose with Particle Swarm Optimization

The neural network predictions of body and wing pose were close to the actual wing pose as observed in the images, but not accurate enough to study the influence of muscle activity and resulting aerodynamic forces, which are extraordinarily sensitive to small changes in kinematics[3]. A larger training set might have resolved this issue, but many additional frames would likely have been required. Instead, we opted to refine the pose estimate in a second step using Particle Swarm Optimization (PSO)[4]. This approach avoids an extensive manual annotation process while still maintaining the benefits of time-independent tracking.

To implement the refinement step, it was necessary to segment the images and create binary masks of the body and wing pixels (Extended Data Fig. 2a). Instead of using a median image for background subtraction, we determined a minimum pixel intensity image over a batch of 100 frames. Body pixels were found by a simple threshold (intensity>200) of the raw, 8-bit image data. Wing pixels were identified by subtracting the minimum image from the frame and subsequently selecting all pixels that had an intensity >10. Subtracting the minimum image removed all stationary pixels, thereby identifying those corresponding to moving body parts.

Our refinement approach used area matching as a measure of how well the 3D model aligned with the images. The advantage of area matching is that it is simple and computationally efficient; however, the disadvantages are that the cost function is not continuous and gradient-based optimization algorithms will not work well. Fortunately, PSO is a gradient-free optimization algorithm that has the ability to find the global minimum, even when the cost function contains multiple local minima. PSO is not guaranteed to converge on the global minimum in a finite number of iterations, but for complex problems such as area matching, it is one of the few optimization methods available.

PSO relies on a swarm of particles that move through the state space with a position and a velocity. The state vector consists of all variables that affect the cost function. At the start of the PSO algorithm, the particles in the swarm are given random positions and random velocity vectors. For a set number of iterations, the algorithm updates the position and velocity of each particle according to the following equations:

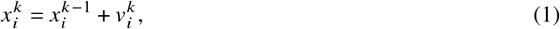

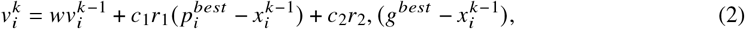

where *i* is the particle index, 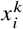 is the particle position at iteration *k*, 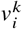 is the particle velocity, *r*_1_ and *r*_2_ are random weight vectors drawn from a normal distribution, 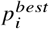 is the personal best of the particle, *g*^*best*^ is the global best for the whole particle swarm, and *w, c*_1_, and *c*_2_ are weights. A particle’s personal best is the position with the lowest value for the cost function throughout the travel history. The global best is the position with minimum cost value in the travel history of all particles of the swarm. In each iteration, the cost function is evaluated for the current position of each particle in the swarm. Subsequently, the cost function values of the personal and global best are compared to the cost evaluations for the current positions. If the current cost function of a particle position is lower than the personal or global best, these values and the associated position vectors are updated. Additionally, in each iteration vectors *r*_1_ and *r*_2_ are updated by randomized weights drawn from a normal distribution.

The velocity update of each particle (equation 2) consists of three parts. The inertia term, *w* 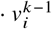 is the previous velocity multiplied with weight, *w*. The second term points towards the personal best of the particle with some added random noise and multiplied by the scaling factor, *c*_1_. The third term is a vector pointing towards the global best of the swarm with added random noise multiplied by a scaling factor, *c*_2_. This scaled randomization introduces stochasticity in the search process, thus insuring that the particles will sample the space near a minimum more often. The vectors pointing towards the personal and global best positions help the particles converge towards a minimum value. The inertia term controls how quickly the swarm converges to a global best. If the inertia term is small, the swarm will converge quickly and risks getting stuck in a local minimum. Setting the inertia term too high, however, may result in no convergence at all or convergence only after a large number of iterations.

In our particular application of PSO, the cost function was based on the projected images of the 3D model. In order to speed up evaluation, the PSO algorithm and the cost function were implemented in C++. The PSO algorithm lends itself well for parallel processing, as the cost function evaluations of all particles can be executed at the same time. Using the std-library in C++17, a separate thread for evaluating the cost function was assigned to each particle. The cost function relies on matrix operations that were implemented using the Armadillo library. To integrate the C++ code within the Flynet GUI, the Boost library was used to make the functions callable in Python.

The 37-dimensions pose vector was a large parameter space in which to search. However, we simplified the process by splitting the pose vector into 5 different components (head, thorax, abdomen, left wing, and right wing), with 7 or 8 state parameters each. Although the pose vectors of the five model components are independent, the cost function evaluation is not, as different model components can overlap in the projected view of the 3D model. To address this dependence, the PSO algorithm updated one randomly selected component at a time.

The variables in the pose vector are all linear, except for the quaternion. A unit quaternion, ∥*q*∥ = 1, can be seen as a point on a 4D sphere with radius 1. However, when updating the quaternion according to equations 1 and 2, the result is not likely to be a unit quaternion. Determining the quaternion difference that satisfies the constraint of a 4D sphere with radius 1 requires application of the concepts of quaternion multiplication and the quaternion conjugate[5]. The details by which we implemented the PSO algorithm using quaternion operations are detailed elsewhere[6].

The cost function for each model component *j*, *C* _*j*_, is given by:

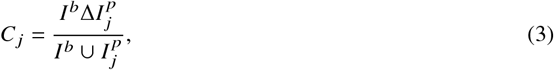

for body components, and

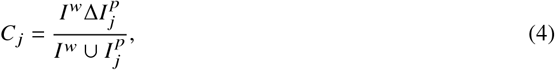

for the wings, where *I*^*b*^, and *I* ^*w*^ are the dynamic bitsets of the body and wing pixels respectively, 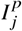 is the dynamic bitset of the model component projections in all 3 views, and Δ and ∪ are the bitwise symmetric difference and union operators, respectively. We vectorized both the images masks and the projected images to accelerate these operations.

Before the tracking started, we set the number of particles, the number of iterations, and the standard deviation of the normal distribution that is used to perturb the initial state and search parameters. The initial particle pose vectors were created by adding random noise to the predicted pose vector from the CNN. Similarly, the particle velocities were randomly sampled from a normal distribution. By specifying the standard deviation of the normal distribution used to generate the random noise vectors, we specified how close the particles searched around the initial pose. We chose a standard deviation of 0.1, which meant that most particles searched around the initial pose value.

With 300 iterations and 16 or more particles, the PSO algorithm converged on the manually annotated pose if the initial CNN prediction was sufficiently accurate. After refining wing pose using PSO, we smoothed the data using an extended Kalman filter as described in detail elsewhere[6].

### Representing wing motion in the strokeplane reference frame

Although mathematically preferable, quaternions do not provide an intuitive sense of wing motion. Instead, we defined a set of Tait-Bryan angles relative to a thorax-fixed reference frame. Prior studies of free flight kinematics[3, 7, 8] have used this approach to define a reference frame at a fixed angle relative to the longitudinal body axis. In case of tethered flight, however, the animal often deflects its head and abdomen for prolonged periods as it attempts to fictively steer. This is problematic, as the body’s longitudinal axis does not necessarily align with the symmetry plane of the thorax under these conditions. Using the pose vector of only the thorax is problematic because its quaternion is not sufficiently accurate. Instead of determining a reference axis based on the body, we defined our wing reference frame using the positions of the left and right wing roots. Because the fly is tethered, the wing roots move in a sable, ’c’-shaped trajectory around the thorax (Fig. 2b). Performing a principle components analysis (PCA) on the left and right root traces provides a convenient means of defining a strokeplane reference frame (SRF). The first three principal components correspond to the *y*-, *x*-, and *z*-axis of the SRF, respectively. Although the axes defined in this way are orthogonal, they are not fixed consistently in space. To create a reference frame that has the *x*-axis pointing forward and the *z*-axis pointing upward, we used the thorax pose vector to establish directionality. Using the normalized axes of the SRF, it is possible to compute the quaternion of the SRF, *q*_*SRF*_. The left and right wing quaternions relative to the SRF may be obtained by multiplying the left and right wing quaternions with the conjugate, 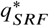. The left and right wing root traces can then be expressed relative to the SRF by subtracting the origin location of the SRF and multiplying by the rotation matrix of 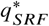

The mechanics of the wing hinge are complex and the root of the wing is not realistically approximated as a single point. This is why we choose not to define any constraints between the wing and thorax in Flynet. However, to simplify the analysis of wing motion and aerodynamics, we assumed the hinge to act as a ball joint at a fixed position on the thorax with 3 degrees of freedom. From an aerodynamics perspective, this assumption is not likely to change computed or measured aerodynamic forces substantially, as the arm of the wing root is relatively small and does not increase wing velocity significantly.

The three Tait-Bryan angles that specify wing orientation are: 1) the stroke angle, *ϕ*, that describes the angle between the *y*-axis of the SRF and the projection of the wingtip on the *x*-*y* plane, 2) the deviation angle, *θ*, that is the angle between the wingtip and the *x*-*y* plane, and 3) the wing pitch angle, *η*, that specifies the orientation of the leading edge panel of the wing with respect to the *z*-axis (Fig. 2c). Wing shape is described by *ξ* and remains unaltered from the Flynet definition. The wing kinematic angles of the right wing are similarly defined such that the signs are symmetric with respect to the left wing.

With the wing kinematic angles defined, we parsed the left and right wing motion into distinct wingbeats. A wingbeat was defined as the time between two subsequent dorsal stroke reversals. We computed the derivative of the stroke angle, *∂ϕ*/*∂t*, and used the condition *ϕ* > 0 and *∂ϕ*/*∂* > 0 to find the dorsal stroke reversal times of the left and right wings in each video sequence. The dorsal reversal time of the left wing does not necessarily align with the right wing, and can be up to 3 frames apart. To remedy these small phases differences, we computed an average time point between the dorsal reversals of the two wings and subsequently rounded to the closest time point. For each high-speed video video sequence, the average dorsal stroke reversal times were determined and used to parse out individual wingbeats, which we indexed so as to preserve the instantaneous frequency and the numerical position within each sequence.

### Encoding wing motion with Legendre polynomials

Depending on the activation level of the indirect power and tension muscles, wingbeat frequencies can vary between 150 Hz and 250 Hz[9]. This variation in wingbeat duration makes it difficult to compare wing kinematics between different high-speed video sequences. A wingbeat frequency of 200 Hz corresponds to about 75 high-speed video frames when captured at 15,000 frames per second. As there are four kinematic angles per wing, there are thus 300 data points representing wing motion during one wingbeat. In order to reduce the number of data points per wingbeat and allow for comparison across wingbeats, we used Legendre polynomials to parameterize wing kinematic traces. Legendre polynomials are well suited for encoding wing kinematics, as they can fit non-periodic traces and asymmetric waveforms. Fourier coefficients, which are often used for fitting time series, require periodic boundary conditions and are biased to symmetric wave forms when a low number of harmonics is used. Similar to Fourier series, Legendre polynomials form a complete and orthogonal system. An easy way to generate a Legendre polynomial is using Rodrigues’formula:

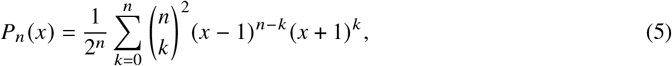

where *n* is the order of the Legendre polynomial and *x* ranges between -1 and 1. A Legendre basis of order *n* is formed by all polynomials from order 0 up to order *n*. By specifying sample points in the range *x* = [−1, 1] and computing the values for each polynomial in the Legendre basis, a Vandermonde matrix is created. The Vandermonde matrix, *X*, can be used in a least-squares fit of a wing kinematic trace, *Y* :

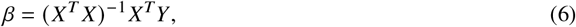

where *β* is a vector of *n*+1 coefficients corresponding to the order of the Legendre basis. With sufficient coefficients it is possible to fit any trace throughout a wingbeat with very little error. When using too many coefficients, however, the the polynomial fit may exhibit high frequency oscillations at the beginning and end of the waveform known as the Runge phenomenon. This may be remedied by lowering the order of the Legendre basis and imposing boundary conditions at the start and end of the wingbeat. Boundary conditions on polynomial fits can be imposed using restricted least-squares[10]. By this method, the restricted least-squares fit, *β*^*\**^, makes use of the the unrestricted least squares fit, *β*, a restriction matrix, *R*, and restriction vector *r*:

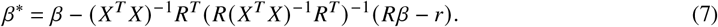

To reduce the Runge phenomenon, we chose the restriction matrix and vectors such that the transition between subsequent wingbeats was continuous up to the fourth derivative. The *j* ^*th*^ derivative of a Legendre polynomial is computed from:

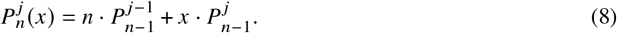

The 1^*st*^, 2^*nd*^, 3^*r d*^, and 4^*th*^ derivatives of the actual wing kinematic trace were computed over a 9 frame time window centered around the start and end time points of the wingbeat. By requiring that the Legendre fit matches the actual values of the wing kinematic trace and the derivatives, the relationship between the least squares fit, *β*^*\**^, the restriction matrix, *R*, and the restriction vector, *r* is:

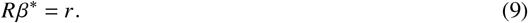

The restriction matrix contains the Legendre basis and the derivative Legendre bases up to the 4^*th*^ derivative for *x* = −1 and *x* = 1 (start and end of the wingbeat). Matching values of the wing kinematic trace and the 1^*st*^, 2^*nd*^, 3^*r d*^, and 4^*th*^ derivatives are contained in the restriction vector *r*. Inserting *R* and *r* in the restricted least-squares solution of equation 7 provides the Legendre coefficients that make the transitions between the previous and next wingbeat continuous up to the 4^*th*^ derivative.

After finding the restricted least-squares solution, we fit the waveforms using higher order Legendre polynomials, making sure to stay below the number of coefficients that induced Runge oscillations. Using higher order polynomials is preferable as it allows for a more accurate fit. For each wing kinematic angle, we tested various number of Legendre polynomials. The stroke angle, *ϕ*, could be fitted accurately with 16 polynomials. The deviation and deformation angles (*θ* and *ξ*) required 20 polynomials each. The wing pitch angle, *η*, required 24 polynomials. Thus, a we used a total of 80 Legendre coefficients to accurately describes the motion of a single wing during each wingbeat.

### Convolutional neural network predicting wing motion from muscle activity

We created a CNN to perform non-linear regression between muscle activity, as measured by GCaMP fluorescence[11], and wing motion, as extracted from the high speed video data using Flynet and PSO. A graphical representation of this code (created using the the model plotting utility in Keras 3.0) is shown in Appendix C; the full code is available as described in the Data Availability Section of the main manuscript.

Before the CNN could be trained, we first normalized the input and output data. To rescale muscle activity traces, the mean, *μ*, and standard deviation, *σ*, were computed for the periods when the fly was flying. A boolean value that specified whether the fly was flying (as determined from wingbeat frequency) was saved to an hdf5 file with a timestamp during each experiment, and could be subsequently used to identify flight bouts and exclude sequences when the animal stopped flying. For all steering muscles, except the *b*_2_ and *iii*_1_ muscles, the activity was rescaled such that a value of 0 corresponds to *μ* − 2*σ* and 1 to *μ* + 2*σ*. The *b*_2_ and *iii*_1_ muscles are typically quiescent during flight, and in some some experiments were not active at all. To accommodate sequences when these muscles were quiescent, the values of *b*_2_ and *iii*_1_ were not rescaled if the standard deviation was below 0.01. If the standard deviation was above this threshold, the *b*_2_ and *iii*_1_ traces were rescaled such that a value of 0 corresponded to *μ* and a value of 1 to *μ* + 3*σ*. Although we tracked wing kinematics for both wings, steering muscle activity was only recorded from the left side of the thorax; thus, we only used the kinematics data from the left wing in our analysis. The 80 values of the Legendre coefficients are in units of radians, which we normalized by dividing by *π*. Wingbeat frequency was rescaled by subtracting 150 Hz from the raw values and subsequently dividing by 100 Hz, such that a value of 0 corresponds to 150 Hz and a value of 1 to 250 Hz.

Although neural networks are capable of handling outliers, too many outlying data points could result in decreased performance. We excluded wingbeats from the dataset under any of the following conditions: 1) the normalized muscle activity was lower than − 0.5 or larger than 1.5, 2) the normalized wingbeat frequency was lower than 0 or larger than 1, and 3) if any of the normalized Legendre coefficients were not inside specified ranges: [−1 /3 <= *C*_*θ*_ <= 1 /3], [−2 3 <= *C*_*η*_ <= 2 /3], [−2 /3 <= *C*_*ϕ*_ <= 2 /3], and [−1 3 <= *C*_*ξ*_ <= 1 /3]. After removing all outliers, the final dataset consisted of 72,219 wingbeats. For the validation set, we selected the first 30 wingbeats of each high-speed video sequence (10,868 total). The remaining wingbeats in each sequence (61,351 total) were used for the training set.

In the first layer of our CNN (Fig. 3), the network processes the GCaMP7f fluorescence signal via a number of convolutional kernels over a fixed time window. After evaluating several different values, we found that a time window length of 9 wingbeats (approximately 45 ms) worked well. A shorter time window did not contain sufficient information and prediction accuracy was worse as a consequence. Longer time windows improved the training error, but the validation error tended to be higher, especially for very long windows (>50 wingbeats).

Our CNN architecture consisted of 2 convolutional layers and 2 dense layers (Fig. 3a). Input of the network consisted of a 13 × 9 matrix of normalized muscle activity and normalized frequency of 9 subsequent wingbeats. The output of the network is a prediction of the 80 normalized Legendre coefficients corresponding to the first wingbeat in a 9 wingbeat time window. Before the input data was fed into the first convolutional layer, Gaussian noise with a standard deviation of 0.05 was added to the input matrix, which helps the CNN to generalize on the data and makes the network more robust to noise in the fluorescence recordings. The first convolutional layer of the CNN consisted of 64 kernels with a 1× 9 kernel window, a1 × 9 stride and SELU activation (Fig. 3a). A total of 256 kernels were used for the second layer, with a 13 × 1 kernel window and stride, and SELU activation. The output of the second layer was a vector with 256 elements. A fully connected dense layer of 1024 virtual neurons takes in the output of the second layer and applies a SELU activation for each virtual neuron. The last layer of the CNN has 80 neurons with a linear activation function, corresponding to the 80 normalized Legendre coefficients.

The CNN was trained for a total of 1000 epochs with a batch size of 100 samples. After every epoch, the network was evaluated using the validation set. The learning rate was set as 10^−4^ with a decay of 10^−7^ per epoch with MSE as a loss function. After the first 200 epochs, the validation error stabilized at approximately 10^−3.7^, whereas the training error continued to decline (below 10^−3.33^) after 1000 epochs (Fig. 3d). After training, the prediction performance of the network was tested on all videos in the dataset. Examples of the CNN prediction performance are given in Fig. 3c and Extended Data Fig. 3. For most sequences, the wing motion predicted from muscle activity was within ±2^°^ of the tracked wing motion for each angle. This accuracy was quite remarkable, given the complex waveforms of the wing kinematic angles, as well as the sparse and filtered nature of the input data.

### Virtual experiments using the trained CNN

After developing a CNN to predict wing motion from observed muscle activity, we used it as an *in-silico* model of the hinge to conduct virtual experiments. The wing kinematics predicted by baseline muscle activity was determined by inspecting the the dataset for sequences that contained no significant changes in muscle activity or wing motion, and subsequently averaging data over those sequences. The baseline values of activity for each muscle and wingbeat frequency, kept constant over the 9 wingbeats, were fed into the trained CNN to yield the baseline pattern of wing motion wingbeat (gray traces, Fig. 4). A naive way of using the CNN to predict the action of individual muscles would be to simply vary the activity of each muscle independently, while keeping that of all other muscles constant constant. However, we observed strong correlations among the activity patterns of many of the muscles within our dataset, determined using RANSAC[12] (Extended Data Fig. 4, Extended Data Table 1). Thus, if we performed virtual experiments by independently changing the activity of each muscle, the input patterns would likely reside outside the data subspace on which the CNN was trained, making the network predictions unreliable. To make more reliable predictions of wing motion from the trained CNN, we deliberately interrogated the CNN using input patterns that would not deviate substantially from those used to train the network by making use of the measured muscle correlations (Extended Data Fig. 4). According to this strategy, we used the trained CNN to probe the influence of each muscle by changing its input value to the maximum normalized value, while simultaneously changing the activity of the other 11 muscles according to the measured correlations.

### Evaluating the function of individual sclerites using latent variable analysis

The CNN that we constructed and trained to predict wing kinematics from muscle activity deliberately mixed input from all 12 control muscles, making it difficult to draw conclusions on the mechanical function of individal wing sclerites. As a complementary approach, we trained a CNN with an encoder-decoder architecture to learn how the activity of all the muscles that attached to a particular sclerite were correlated to changes in wing motion (Fig. 6; Extended Data Fig. 8). The encoder block contained deliberate bottlenecks that forced the network to learn how to represent the input data by a small set of latent variables representing the state of each sclerite and wingbeat frequency. Training the encoder-decoder with a bottleneck is roughly equivalent to performing a non-linear PCA on the dataset[13]. A graphical representation this code (created using the the model plotting utility in Keras 3.0) is shown in Appendix D; the full code is available as described in the Data Availability Section of the main manuscript.

We constructed the encoder-decoder with two decoder heads, one that predicts muscle activity from the latent variable space and another that predicts wing kinematics (Extended Data Fig. 8). Input to the encoder-decoder was split into 5 streams, grouping the activity of muscles that insert on each of the 4 wing sclerites with a separate stream for wingbeat frequency. Each stream uses two convolutional layers, as in our original CNN model (Fig. 3a), but followed by a dense layer of 512 neurons and a final layer of one neuron with linear activation that corresponded to one of the five latent variables. After the latent space, the network is split into two streams: one stream uses a decoder to predict the input to the network (i.e. the muscle activity), the second stream uses two dense layers to predict the Legendre coefficients of the wing motion. We placed a back-propagation stop between the latent space and the muscle activity decoder layers, such that the latent space is only trained on correlations with wing motion. The muscle activity decoder layers are still functional, as the decoder predicts the correlation between the latent space and muscle activity, but without the noise of the input to the network.

We trained the network on the muscle activity and wing motion dataset using 1000 epochs, a batch size of 100, a learning rate of 10^−4^, and a decay rate of 10^−7^. The network predicts two vectors: the latent space (5 × 1) and Legendre coefficients (80 × 1), and one matrix: muscle activity and wingbeat frequency over 9 wingbeats (13 × 9). It is important to note that the prediction error of the wing kinematics by the sclerite function encoder-decoder is expected to be worse than our original CNN, as the bottleneck required to create the latent variable space greatly restricts the amount of information that can be used to learn the correlations. In Figure 6 and Extended Data Fig. 8, only the decoder sections of the network were used to predict muscle activity and wing motion for each latent variable, which we varied systematically between − 3*σ* and + 3*σ* in 9 steps while keeping the other latent variables constant at 0. The muscle activity and wing motion were then predicted from the latent input by the two decoders.

### Dynamically-scaled flapping wing experiments

In the last three decades, our laboratory has developed several versions of a dynamically-scaled flapping wing robot[14–18]; however, none of these had the capability of creating the chord-wise deformations that we observed and tracked in our high-speed videos. The robotic wings we designed for this study were actuated by one stepper motor and two servo motors each (Extended Data Fig. 5a). Stroke angle was controlled by a stepper motor (10 kHz clock) via two gears with a 1:3 gear ratio. The position of the stepper motor was controlled via micro-stepping, which permitted fine motor control (7.8·10^−3^ degrees per microstep). Two magnets were positioned on the gear so that a Hall-effect sensor was activated when the wing moved out of bounds. Besides protecting the wing, the Hall-effect sensor was also used to home the stepper motor. During an experiment, a Teensy 3.2 microcontroller tracked the number of microsteps travelled relative to the home position. To account for motor slip, which sometimes occurred due to large torques, the steppers were homed after each experiment. During an experiment, the stepper’s position, *ϕ*, and velocity, 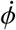, were controlled via a feed-forward process. The stepper made micro-steps on a 10 kHz clock; position and velocity set points were updated at 200 Hz. The custom instrumentation software computed the required stepping frequency and direction to reach the specified position and velocity values after each update.

The deviation and wing pitch angles were controlled by two servos (HiTec D951TW). The deviation angle servo operated via two gears (1:1 gear ratio), and the wing was directly attached by the wing pitch servo. The position of the servos was specified by a pulse-width modulation (PWM) signal at 50 Hz. The deviation servo could move between −45^°^ and +45^°^ and the wing pitch servo could move between +90^°^ and −90^°^. During an experiment, the position of the servo was updated at 50 Hz in a feed-forward control loop.

We measured forces and torques on the wing using a 6 degree-of-freedom force and torque (FT) sensor (ATI Nano 17), mounted on the rotation axis of the wing pitch servo. Custom-machined aluminium mounts coupled the base of the FT sensor to the servo axis and the wing. During each experiment, six 16-bit unsigned integers corresponding to all forces and torques (*F*_*x*_, *F*_*y*_, *F*_*z*_, *T*_*x*_, *T*_*y*_, *T*_*z*_) were sampled at 200 Hz. The robot was submerged in an acrylic tank (2.4× 1× 1.2 *m*) filled with mineral oil (Chevron Superla white oil), with a kinematic viscosity of 115 · 10^−6^ *m*^2^·*s*^−1^ and a density of 880 *kg* ·*m*^−3^ at 22 *C*^°^[19]. The oil level in the robofly tank was approximately 1 m. The robot had the wingtip radius of 31 cm and was submerged at a depth of 50 cm. A previous study that systematically mapped the effects of the tank boundaries on lift an drag determined that the system approximated the forces generated within an infinite volume, provided the wing remains further than 12 cm from any boundary (top, bottom, or side)[14].

Besides the three Tait-Bryan angles describing wing orientation, Flynet tracked a fourth angle describing wing shape, the deformation angle *ξ*. To implement this fourth angle on the robot, the wing was composed of four panels connected by three hinge lines (Exended Data Fig. 6a). The four panels were cut out of an acrylic sheet (2.75 mm thickness). Each hinge consisted of a 2 mm steel rod core, surrounded by an acrylic tube with inner and outer diameters of 2 mm and 4 mm, respectively. The acrylic tube was cut into sections of 20 mm and these sections were glued in an alternating pattern to two adjacent panels. The rotation angle between two subsequent panels was controlled by a micro servo (HiTec HS-7115TH), which was screwed onto one panel and connected to the next panel via a 1 mm metal rod that was coupled to the servo arm. Three micro servos were used to deform the wing. Following the assumption that the wing bending angle was uniformly distributed over the three hinge lines, the deformation angle *ξ* was divided by 3 to obtain the rotation angle per hinge line.

Dynamic scaling was ensured by matching the Reynolds number of the robotic wing and a real fly, which requires:

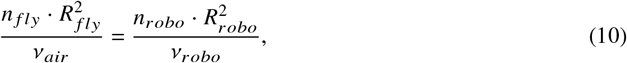

where *n* _*fly*_ is the fly’s wingbeat frequency, *n*_*robo*_ the wingbeat frequency of the robot, *R* _*fly*_ the fly’s wing length, *R*_*robo*_ the wing length of the robot, *v*_*air*_ the kinematic viscosity of air and *v*_*robo*_ the kinematic viscosity of the mineral oil. Rewriting the Reynolds number equality yields:

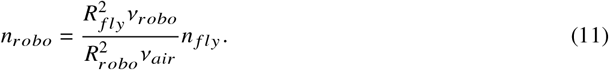

By entering the values for a typical fly (*R* _*fly*_ = 2.7*mm, v*_*air*_ = 15.7 ·10^−6^*m*^2^*s*^−1^ at 25^°^*C, n* _*fly*_ = 200*Hz*) and the RoboFly parameters (*R*_*robo*_ = 310*mm, v*_*robo*_ = 115· 10^−6^*m*^2^*s*^−1^ at 22^°^*C*), the required flapping frequency for the robot is 0.11*Hz*. The force scaling factor may be written as[6]:

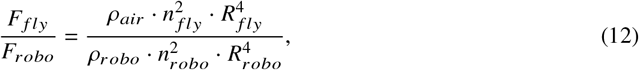

with relevant density values of *ρ*_*air*_ = 1.18*kgm*^−3^ and *ρ*_*robo*_ = 880*kgm*^−3^. The torque scaling factor is the product of force and moment arm:

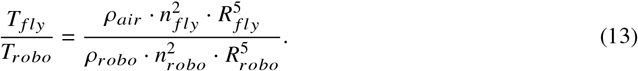

Any experiment using the robotic wing required an input file that specified the following angles: ϕ,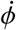, *θ, η* and ξ at intervals of 5 ms. A C++ program read the five angles and sent them to the Teensy microcontroller via a USB-cable at 200 Hz (RawHID protocol). The Teensy microcontroller converted the angles to PWM commands for the servos and step&direction commands for the stepper motors. At the same time, the force and torque data were sent from the Teensy micro-controller to the C++ program, which logged the measurements.

At the start of an experiment, the wing was moved to the home position (ϕ = 0, *θ* = 0, *η* = 0, ξ = 0). In order to prevent rapid acceleration of the wing, the wing kinematic angles during the first wingbeat of an experiment were multiplied with the following function:

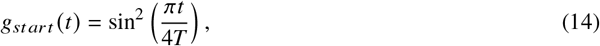

where *T* corresponds to the wingbeat period. Similarly, the last wingbeat of the experiment ends at the home position and the wing kinematic angles are multiplied by:

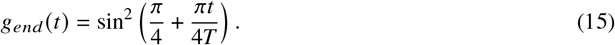

When wings start flapping in the oil, it takes approximately two to three wingbeats before the wake is fully developed[15]. In order to exclude these start-up effects, a wing kinematic pattern was repeated for 9 wingbeats and only wingbeats 4 through 8 were used for analysis. After each experiment, the data were converted from 16-bit unsigned integers into actual force and torque vectors and smoothed using a linear Kalman filter. Besides aerodynamic forces, the FT-sensor also measured inertial and gravitational components. The low flapping frequency of the robot assures that the inertial forces may be ignored; however, the gravitational forces are significant. To remove the gravitational force from the analysis, every wing kinematic pattern was replayed at a 5x slower frequency, which reduces aerodynamic forces by a factor of 25. The forces and moment measured during the slow frequency, which approximate the gravity component, were subtracted from the fast frequency data after interpolation.

The aerodynamic and gravitational forces were measured by the FT sensor in the wing reference frame. For subsequent analysis it is useful to translate the forces and torques to the stroke reference frame (SRF), which was accomplished by the following Tait-Bryan operations:

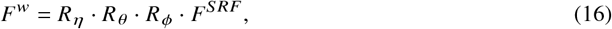

and

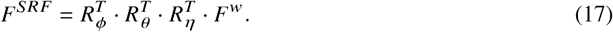

where *F*^*w*^ and *F*^*SRF*^ are the forces in the two different reference frames, and *F*^*w*^ and *R*_*η*_, *R*_*θ*_, *R*_*η*_, correspond to the rotation matrices. More details on thse operations are provided elsewhere[6].

### Computing inertial forces via the Newton-Euler equations

Besides aerodynamic forces, wing inertia plays a significant role in fly flight. Although the wing mass is only ∼0.2% of the body mass[9], the angular velocity and acceleration of wing motion is very high. We estimated the inertial forces and torques of a rotating rigid body using the Newton-Euler equations:

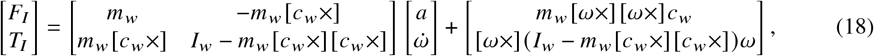

where *m*_*w*_ corresponds to the wing mass, *c*_*w*_ is the position of the center of gravity on the wing, *I*_*w*_ is the inertia tensor of the wing, *a* is the linear acceleration of the wing, 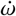 is the angular acceleration, and *ω* the angular velocity, with all values relative to the wing reference frame. We ignored the effects of the linear acceleration of the wing because they are very small compared to the angular acceleration. The first right-hand term in equation 18 corresponds to the inertial forces and torques due to wing acceleration, *F* _*acc*_ and *T* _*acc*_. Inertial effects dependent on angular velocity, such as the Coriolis and the centrifugal forces, are captured by the second right-hand term and are collectively referred to as *F*_*angvel*_ and *T*_*angvel*_. We assumed that the inertial effects of wing deformation are small, and thus calculated inertial forces and torques based on the assumption that the wing is a rigid, flat plate.

The angular velocity and acceleration had to be computed from the Legendre polynomials describing wing motion. We computed the emporal derivatives of the Legendre polynomials 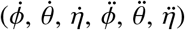 using equation 8. However, the temporal derivatives of the wing kinematic angles do not correspond to the angular velocity and acceleration. Angular velocity in the wing reference frame is given by:

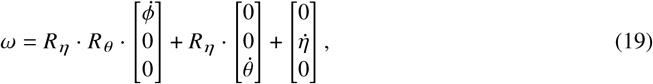

where *R*_*θ*_ and *R*_*η*_ correspond to the rotation matrices of the Tait-Bryan operations for *θ* and *η*, respectively. Angular acceleration is given by:

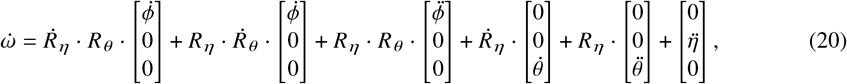

with 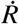 being the temporal derivative of the rotation matrix.

To compute the inertial forces and torques, the mass, center of gravity, and the inertia tensor of the wing must be determined. These parameters were computed using the scaled 3D model we created for Flynet, with a wing length of 2.7*mm*, an estimated cuticle density of 1200*kg*·*m*^−3^, and a wing thickness of 5.4*μm*[20]. The inertial parameters of the (left) wing are given by:

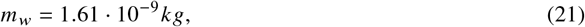

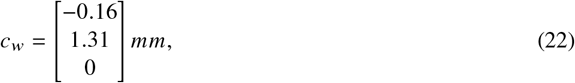

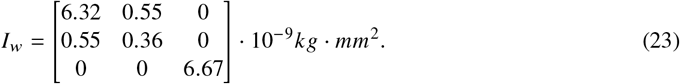

With the wing inertia parameters, angular velocity, and acceleration defined, it is possible to compute the inertial forces and torques for any arbitrary pattern of wing motion pattern. Extended Data Fig. 5b shows the aerodynamic and inertial forces and torques in the SRF for the baseline wingbeat. The aerodynamic forces were measured on the robot and rescaled to the appropriate units for an actual fly using equations 12 and 13. To make interpretation easier, the forces and torques have been non-dimensionalized by dividing the forces by the body weight, and the torques by product of body weight and wing length. Using the estimated cuticle density and the scaled body components of the 3D fly model, the body mass was found to be 1.16 ·10^−6^ *kg*. Inspection of the traces in Extended Data Fig. 5b indicate that the inertial forces and torques in wing motion are substantial, as found in prior studies[21].

With the methods to measure aerodynamic forces and compute inertial forces established, it is possible to evaluate the forces generated by changes in steering muscle activity. For each steering muscle, seven wing kinematic patterns were tested on the robotic wing. The seven wing kinematic patterns were found by sampling the muscle activity patterns on a line between the baseline muscle activity and the maximum muscle activity of a selected muscle (Fig. 4), and subsequently predicting the corresponding wing motion using the trained CNN. The workflow to obtain the control forces and torques of muscle activity consisted of measuring the force and torque traces of 84 wing kinematic patterns on the robot, performing the gravity subtraction, smoothing the traces with a Kalman filter, converting traces to the SRF, computing the median force and torque traces from the measured wingbeats (*F* _*aero*_ and *T* _*aero*_), estimating the inertial components (*F* _*acc*_, *T* _*acc*_,*F* _*angvel*_, *T* _*angvel*_), and finally, summing the inertial and aerodynamic forces and torques (*F*_*total*_ and *T* _*total*_). A time history of the individual force and torque components are plotted over a baseline wingbeat in Extended Data Fig. 5b. The corresponding instantaneous total force vectors are shown superimposed over the 3D kinematics in Extended Data Fig. 5c, with comparable plots for the kinematics in each virtual muscle activation experiment (Fig. 4) shown in Extended Data Fig 6. Further details on the construction and operation of the robot and the analysis of aerodynamic forces and torques are provided elsewhere[6]

### Simulating arbitrary free flight maneuvers with Model Predictive Control

Using our CNN, the virtual muscle activation experiments, and robotic wing experiments provides us with the components necessary to construct a state-space simulation of a flying fly using Model Predictive Control (MPC)[22], in which the virtual fly regulates its flight trajectory via changes in the time course of steering muscle activation. By specifying an initial and goal state and a time period to achieve the goal state, a MPC controller finds the optimal trajectory given a cost function and constraints. The explicit discrete time-invariant state-space system is governed by the following equations:

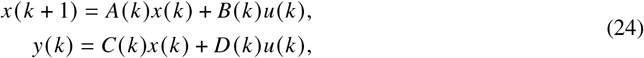

where *x*(*k*) and *x* (*k*+ 1) are the state vectors at times *k* and *k*+ 1 respectively, *u*(*k*) is the control vector, *A* (*k*) is the system matrix, *B* (*k*) is the control matrix, *C* (*k*) the output matrix, *D* (*k*) the feed-forward matrix, and *y* (*k*) the output vector. In the case of fly flight, there is no feed-forward process and the output of the state-space system is the state vector. This means that matrices *C* (*k*) and *D* (*k*) do not need to be defined (Extended Data Fig. 7a).

The state vector describes the orientation, position, velocity, and angular velocity of the fly’s body:

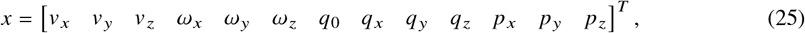

where *v* and *ω* are the linear and angular velocity of the body in the SRF, respectively, *q* is the quaternion describing SRF orientation relative to the inertial reference frame, and *p* is the position of the SRF in the inertial reference frame. For simplicity, we assume that the head and abdomen are stationary relative to the thorax. The temporal derivative of the state vector, 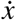, is required for updating the state for each time step:

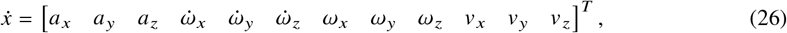

where *a* and 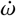 are the linear and angular acceleration of the body in the SRF, respectively.

At each time step, the state vector is multiplied with the system matrix, *A* (*k*), to compute the temporal derivative of the state vector. The time step, Δ*t*, corresponds to the wingbeat period, which is assumed to be constant at 1/*f* = 1/200 = 0.005 *s*. Although flies regulate wingbeat frequency during flight, it typically takes 20 wingbeats to increase or decrease the wingbeat frequency to a new level[3]. As our model was developed for simulating rapid maneuvers of only 10 wingbeats, frequency was held constant in our simulations.

The equations of motion of the system matrix include the following flight forces and torques: body weight, body inertia, body aerodynamics, and the aerodynamic and inertial damping of the wings. The mass, center of gravity, and inertia tensor of the body were determined from the scaled 3D body model (head, thorax, abdomen) for the average fly in the dataset. For the wing mass, the cuticle density was assumed to be 1200 *kg* · *m*^−3^, and the body mass estimated as *m*_*b*_ = 1.16 · 10^−6^ *kg*. The center of gravity of the body in the SRF was estimated as *cb*=[0.04 0 −0.24] ^*T*^ *mm*. The inertia tensor of the body is given by:

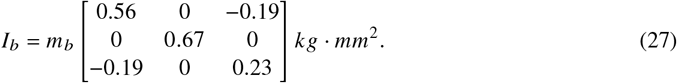

The inertial forces and torques on the body can be computed with the Newton-Euler equations, evaluated at the center of gravity of the body:

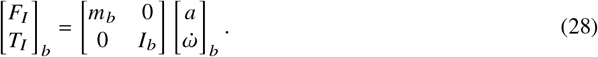

Obtaining the rotational accelerations of the body requires solving the inverse problem:

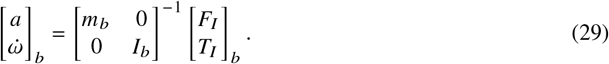

As the state is evaluated at the center of gravity, there are no torques due to the fly’s weight. Although the aerodynamics of the wings generate larger forces, the body of the fly experiences drag during flight. The combined 3D shape of the head, thorax, abdomen and legs is complex, but we chose to model the body conservatively as sphere with a radius of 1 *mm*. While the actual body drag is likely to be lower, the simplicity of the drag model makes it easier to implement in the state-space system. The drag on the sphere can be calculated as:

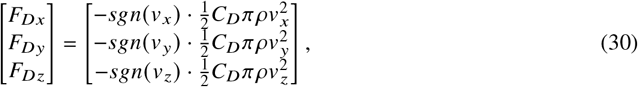

with the drag coefficient as *C*_*D*_ = 0.5 and *sgn* as the sign operator. Because the center of pressure of the sphere is assumed to be at the center of gravity of the body, there are no torques due to body drag.

Because the aerodynamic and inertial damping terms due to the motion of the flapping wings are dependent on body rotational velocity, these components are added to the system matrix. The equations of motion to compute the relevant linear and rotational accelerations can be written as:

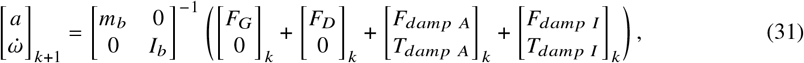

where *F*_*dampA*_ and *T*_*dampA*_ are the aerodynamic damping terms and *F*_*dampI*_, and *T*_*dampI*_ are the inertial damping terms. *F*_*G*_ is the gravity component, expressed in the SRF[6].

The control inputs that are available to the fly are the muscle activity patterns of the left and right steering muscles: *u*^*L*^ and *u*^*R*^. The sum of aerodynamic and inertial forces and torques generated via changes in these inputs can be computed as:

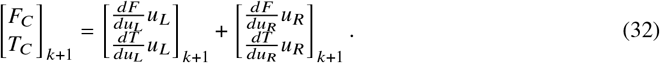

Adding these control forces and torques to equation 31 gives the complete equations of motion for body rotational acceleration:

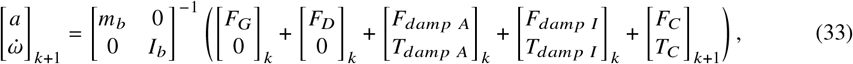

The computed accelerations of equation 33 are subsequently be used to update the state vector. As the body accelerations and velocities are both in the SRF, the update of the body velocity is relatively simple:

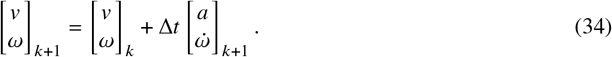

The body quaternion and position are in the inertial reference frame and the angular velocity update must be transformed from the SRF to the inertial reference frame. Using quaternion multiplication, the quaternion update can be written as:

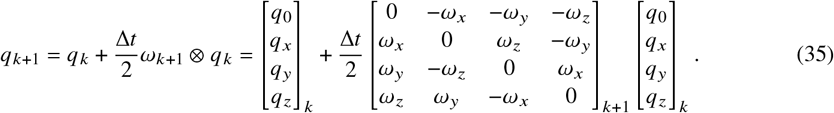

After computing *q*_*k*+1_, the quaternion must be normalized such that ∥*q*∥ = 1. The position update is given by:

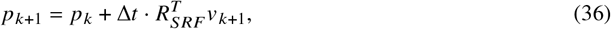

where 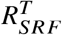 is computed using *q*_*k*_.

The aerodynamic and inertial forces for the maximum muscle activity patterns were measured and computed in a stationary reference frame. During free flight, the body translates and rotates, which has an effect on both the inertial and aerodynamic forces on the wing[23–25]. We used a quasi-steady aerodynamic model and the Newton-Euler equations to compute the relevant aerodynamic and inertial damping coefficients for fruit fly flight. The effects of inertial damping, can be computed given the angular velocity of the wing:

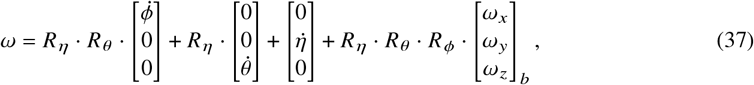

where the last term adds the body angular velocity, converted to the wing reference frame. By inserting the redefined angular velocities of the left and right wings into the Newton-Euler equations, it is possible to compute the inertial forces and torques given a constant body angular velocity.

The wingbeat-averaged inertial forces and torques that are generated by body rotation show a linear trend with angular velocity. Translational body velocity does not cause any changes in inertial force or torque. The slopes of the linear trends form the inertial damping matrix:

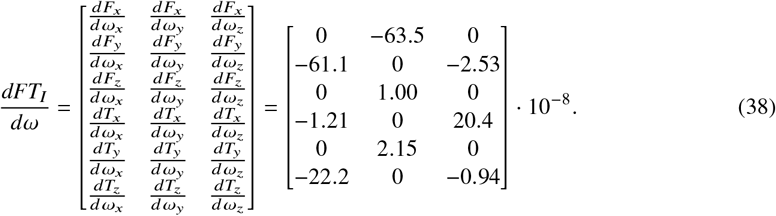

Rotational body velocity generates a torque in opposite direction for *ω*_*x*_ and *ω*_*z*_, but a torque in the same direction for *ω*_*y*_.The positive damping torque for pitch rotation means that flight is unstable around the pitch axis. There is a strong coupling between the roll rotation and yaw torque, and vice versa. Aerodynamic damping was computed using a quasi-steady model, the details of which are described elsewhere[6].

The angular velocity of the body can be added to the angular velocity of the wing using equation 37. Inclusion of the translational body velocity into the wing’s angular velocity term is difficult, however. We therefore implemented the translational and rotational quasi-steady terms using a blade-element formulation, in which the wing is partitioned into spanwise sections and the air velocity vector is computed on each section[26]. Simulating the baseline wing kinematics on the left and right wings, with the relevant body velocities yields the damping matrix for body translational velocity:

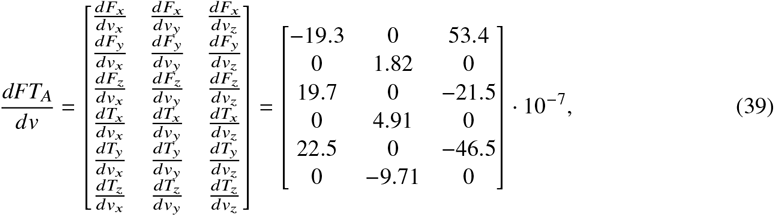

and the damping matrix for body angular velocity:

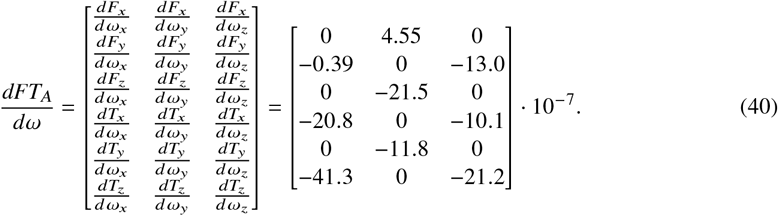

The aerodynamic damping coefficients are approximately an order of magnitude larger than the inertial damping coefficients. Both *dF*_*Ax*_/*dv*_*x*_ and *dF*_*Az*_ /*dv*_*z*_ have a negative sign, which means that there is a force opposite to the direction of body motion. In cases of *dF*_*Ay*_ /*dv*_*y*_, *dF*_*Az*_/*dv*_*x*_, and *dF*_*Ax*_ /*dv*_*z*_, the signs are positive, however; which indicates a force in the direction of body motion. Whether the damping coefficient is negative or positive depends on the effects of body velocity on parameters such as angle-of-attack, instantaneous air velocity, and wing orientation. All the aerodynamic damping torques, *dT*_*A*_ /*dω*, have a negative sign. This means that for the baseline wing motion, flight is stable to perturbations in angular velocity, a conclusion consistent with a prior study[23].Matrices *dFT*_*I*_ *dω, dFT*_*A*_ /*dv*, and *dFT*_*A*_ /*dω* can be converted into a single damping matrix. Multiplication of the damping matrix with the linear and angular velocity vectors of the body yields the damping forces and torques:

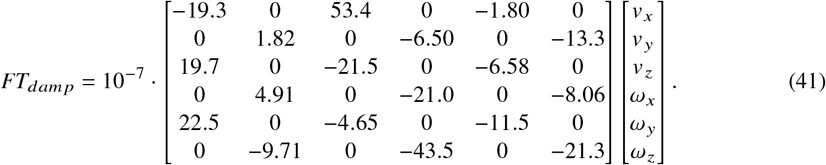

A central feature of our model is that the control inputs available to the fly are the muscle activity patterns of the left and right steering muscles: *u*_*L*_, *u*_*R*_. Using the measurements from the robotic wing, it is possible to compute the Jacobians (*dFT*/*du*_*L*_, *dFT*/*du*_*R*_) of the aerodynamic force and torque production for maximum muscle activity pattern of each muscle. Steering muscle activity is ordered as follows in the control vectors:

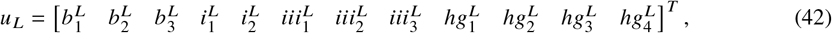

and

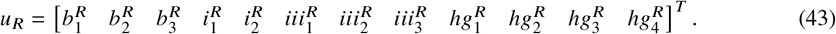

A full table of numerical values of the calculated Jacobians for the aerodynamic force and torque production of the left and right for each steering muscles are provided elsewhere[6]. Control forces and torques can be computed by multiplying the Jacobian with the control vector:

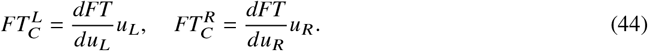

The muscle activity correlation analysis in Extended Data Fig. 4 shows that not all combinations of muscle activity are possible. It is, therefore, necessary to impose constraints on the left and right control vectors. Specifically, this constraint is that muscle activity has to lie on a 12-D surface representing the muscle correlation matrix (Extended Data Table 1). A mathematical formulation for the 12-D surface constraint is:

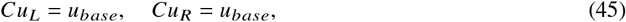

where *C* is the 12 ×12 correlation matrix (Extended Data Table 1), and *u*_*base*_ is the baseline muscle activity pattern.

MPC optimizes trajectories in state-space, over a finite time window, for a given cost function[22]. The state-space system is used to predict a state trajectory over a finite time horizon. At each time step, the controller determines the optimal control input that generates one or more trajectories that minimize the cost function, while being subject to system constraints. This process is repeated for every time step, shifting the horizon forward. MPC is thus also known as receding horizon control.

We used the the Python-package *do-mpc*[27] to implement our MPC simulation. For the free flight simulations, the controller uses discrete time and a non-linear model. The duration of all free flight simulations was 10 wingbeats, and the MPC horizon was set to 10 wingbeats as well. This means that the goal state is always visible to the controller. At the center of MPC is the objective function, which is defined in do-mpc as:

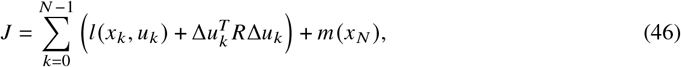

with *N* being the number of time steps of the problem, *l* (*x*_*k*_, *u*_*k*_)is the Lagrange term, 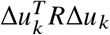 is the r-term, and *m* (*x*_*N*_)is the Meyer term. The Lagrange term provides the option to provide a reference-state trajectory and add a penalty for any deviation. Rapid changes in control input are penalized using the r-term: Δ*u*_*k*_ = *u*_*k*_−*u*_*k*−1_, and *R* is a weighting matrix. The Meyer term evaluates how close the specified goal state is to the final state, *x*_*N*_.

In case of the free flight simulations, the MPC objective function is defined as follows. First, the initial state, *x*_0_, and the goal state, *x*_*n*_, are specified. Subsequently, the Lagrange and Meyer terms are both defined as:

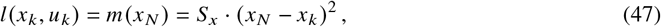

with *S*_*x*_ as a scaling matrix. By setting the diagonal values of the *S*_*x*_ and *R* matrix, the user can tune the importance of certain aspects of the objective function. The Lagrange and Meyer terms we used were always the same, as we did not want to specify any reference trajectory for the Lagrange term. In this way, the objective function allowed the MPC controller to explore any trajectory that ends at the goal state. By setting a diagonal element to zero in the *S*_*x*_ matrix, the MPC controller will not optimize the trajectories for this parameter. If a diagonal element in *S*_*x*_ is set to a high value, the MPC controller will prioritize this element. For example, by setting the weight for the goal, *v*_*y*_, to 1, the controller will punish any deviation from this goal heavily, and allows the user to enforce straight flight. In a similar way, the user can set the diagonal values of the *R* matrix, and determine the behavior of the steering muscles during the simulation. For the free flight simulations, we set all the diagonal values of *R* to 1.

Besides tuning the cost function, the bounds of the state and control vectors must be specified. In case of the control vectors, all steering muscle activity was bounded between 0 and 1. In the state vectors, linear velocity was bounded by − 10^4^ and 10^4^ *mms*^−1^, angular velocity was bounded by -1000 and 1000 *rads*^−1^, quaternion values were bounded by -1 and 1, position was bounded by −10^4^ and 10^4^ *mm*, and the body quaternion was constrained to be a unit quaternion, ∥*q*∥= 1. Do-mpc allows the user to specify the tolerance by which a state or control vector can deviate before a constraint will be enforced. For the free flight simulations, we set the constraint tolerance on the control vector, *c*_*tol*_, to 0.001.

Extended Data Fig. 7a shows the workflow of the MPC controller. The user only needs to specify the initial and goal states, time in which the fly should reach the goal state, and the weight matrix of the objective function, *S*_*x*_. With these parameters, the MPC controller will try to find the optimal state trajectory and required control inputs, that minimizes the objective function while satisfying the constraints. Further details on the design and implementation of our MPC controller may be found elsewhere[6]. Our software is available in the form of a Jupyter notebook as described in the Code Availability Statement associated with our manuscript.

## Appendix A Processes flow for CNN predicting wing pose from high speed images (Flynet, Part 1)

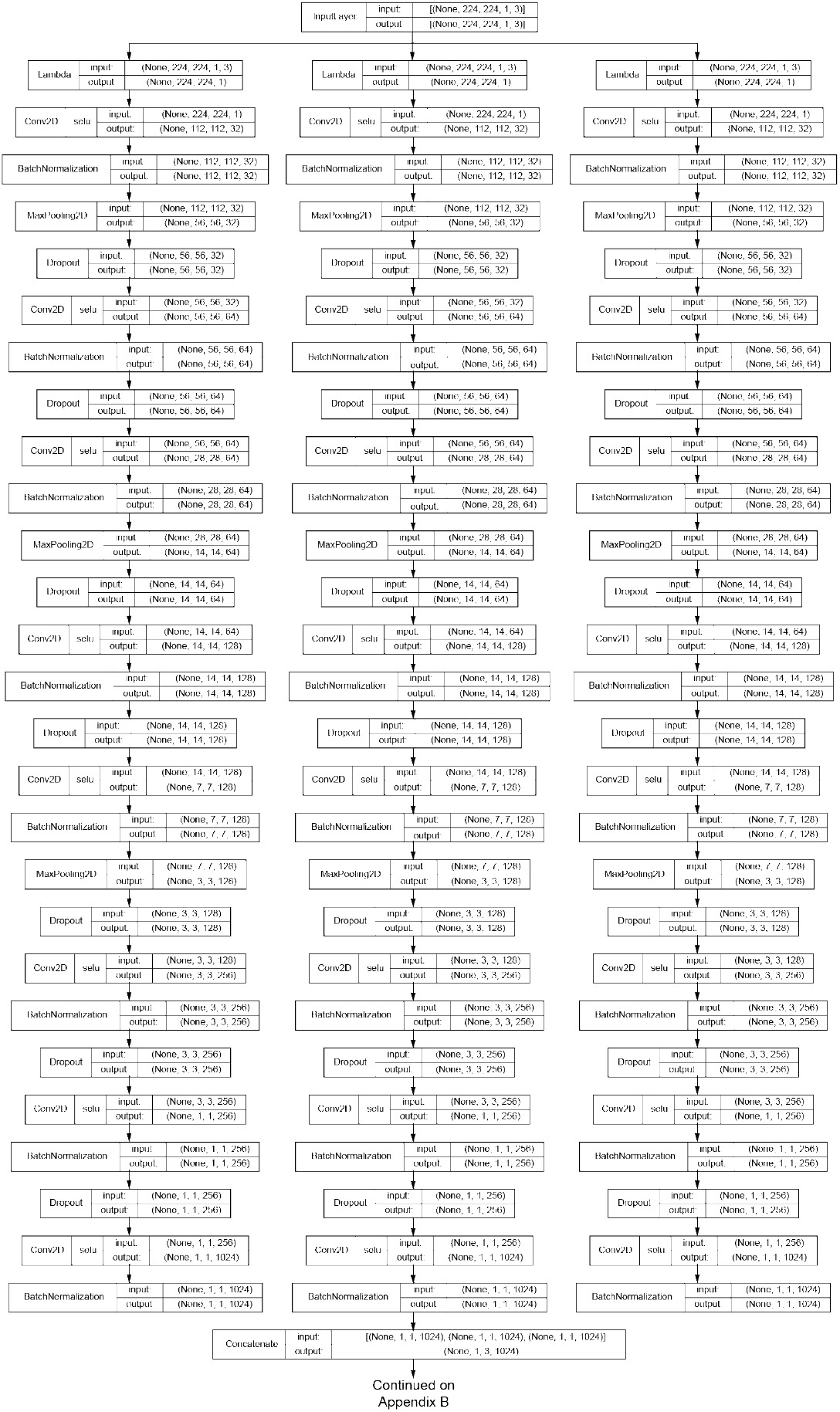

## Appendix B Processes flow for CNN predicting wing pose from high speed images (Flynet, Part 2)

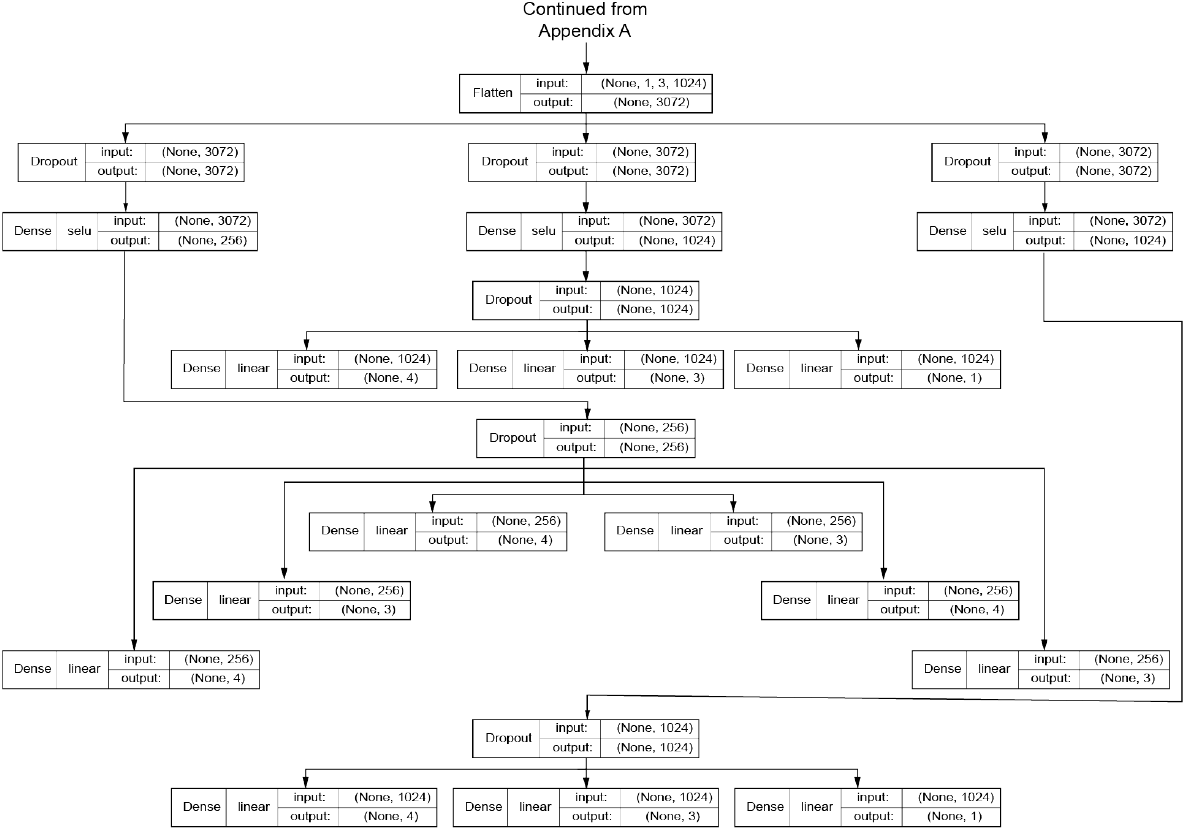

## Appendix C Processes flow for CNN predicting wing kinematics from steering muscle activity

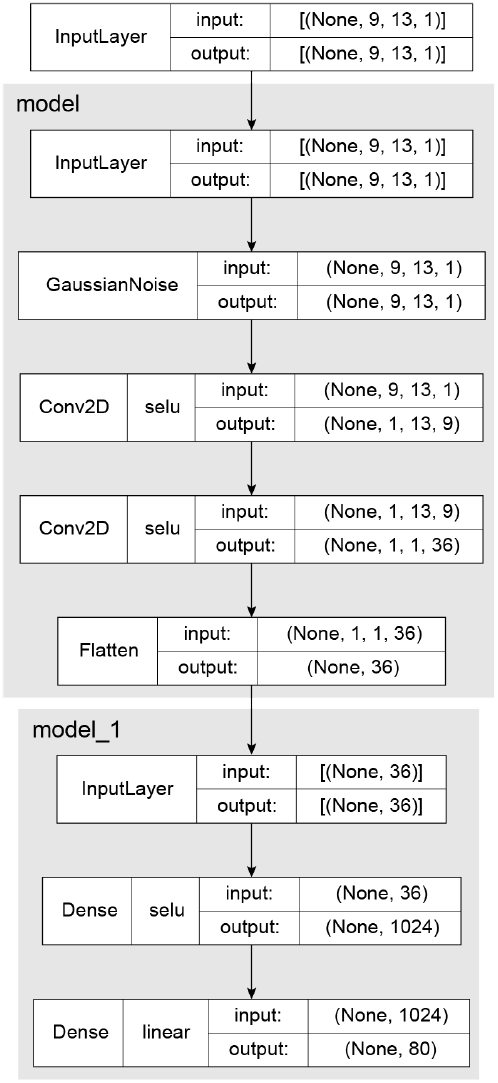

## Appendix D Processes flow for encoder-decoder that performs latent variable analysis

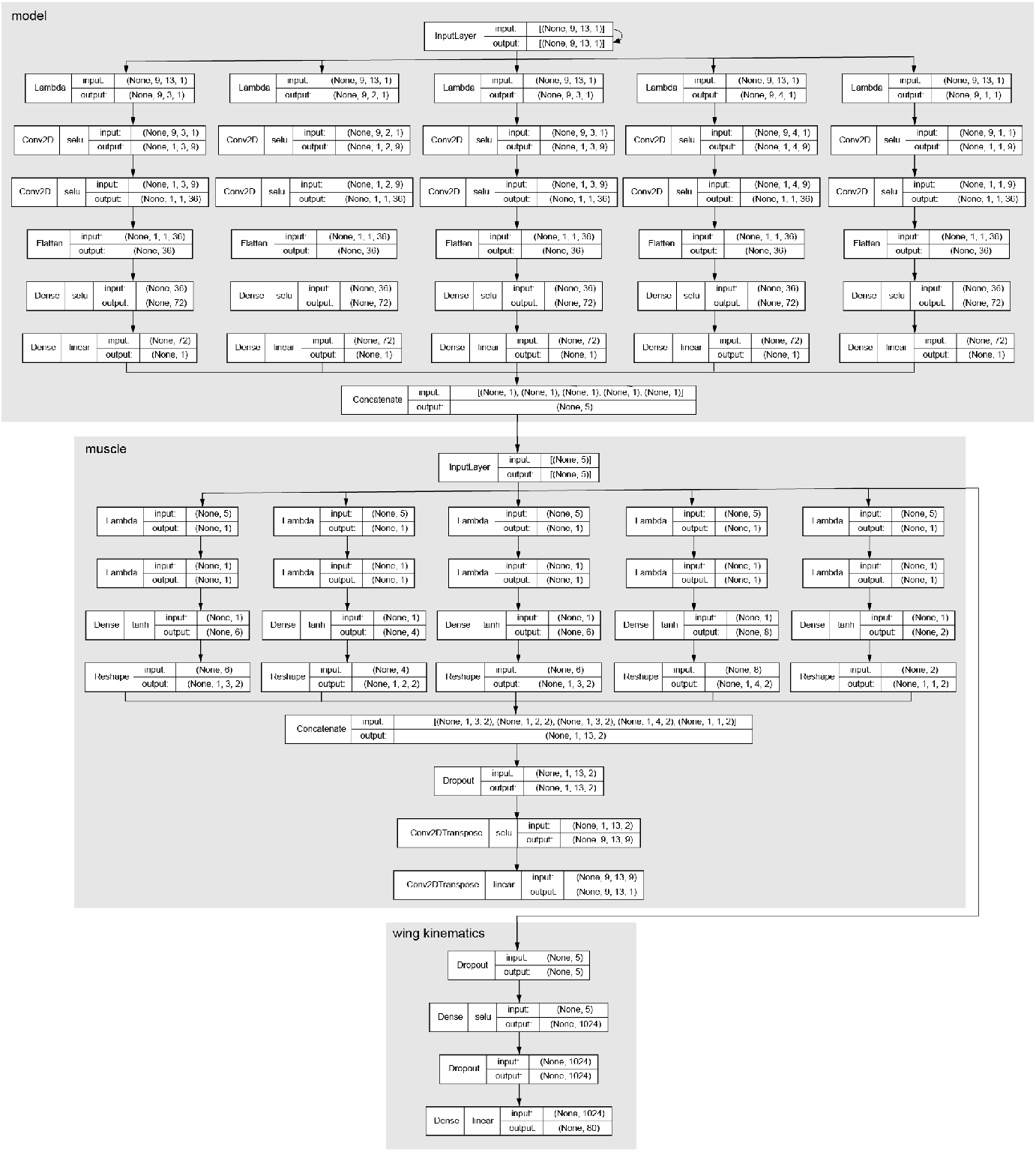

## Notes

### Competing Interest Statement

The authors have declared no competing interest.

### Summary of Updates

Dr. Igor Siwanowicz has been added as an author to this paper. He has provided a detailed morphological analysis of the wing hinge that is presented in a supplementary animation. The revised manuscript contains a more extensive Supplementary Information section that details the mathematical methods used throughout our analysis. The new version also contains a few minor alterations to the text and figures that improve the clarity of the presentation.

